# CO_2_ enhances the ability of *Candida albicans* to form biofilms, overcome nutritional immunity and resist antifungal treatment

**DOI:** 10.1101/2020.03.31.018200

**Authors:** Daniel R. Pentland, Fritz A. Mühlschlegel, Campbell W. Gourlay

## Abstract

*C. albicans* is the predominant fungal pathogen of humans and frequently colonises medical devices, such as voice prosthesis, as a biofilm. It is a dimorphic yeast that can switch between yeast and hyphal forms in response to environmental cues, a property that is essential during biofilm establishment and maturation. One such cue is the elevation of CO_2_ levels, as observed in exhaled breath.. However, despite the clear medical relevance, the effects of CO_2_ on *C. albicans* biofilm growth has not been investigated to date. Here, we show that physiologically relevant CO_2_ elevation enhances each stage of the *C. albicans* biofilm forming process;from attachment through to maturation and dispersion.. The effects of CO_2_ are mediated via the Ras/cAMP/PKA signalling pathway and the central biofilm regulators Efg1, Brg1, Bcr1 and Ndt80. Biofilms grown under elevated CO_2_ conditions also exhibit increased azole resistance, tolerance to nutritional immunity and enhanced glucose uptake to support their rapid growth. These findings suggest that *C. albicans* has evolved to utilise the CO_2_ signal to promote biofilm formation within the host. We investigate the possibility of targeting CO_2_ activated processes and propose 2-Deoxyglucose as a drug that may be repurposed to prevent *C. albicans* biofilm formation on medical airway management implants. We thus characterise the mechanisms by which CO_2_ promotes *C. albicans* biofilm formation and suggest new approaches for future preventative strategies.

## Introduction

*C. albicans* is a commensal yeast located on the mucosal surfaces of the oral cavity, gastrointestinal and genitourinary tracts of most healthy individuals ^12^. Despite being a commensal organism, it is also an opportunistic pathogen ^13^; in fact, it is the most widespread of all the human fungal pathogens ^4^ and is the fourth most common cause of hospital-acquired infections in the USA ^1^. Infection with *C. albicans* is a particular problem among immunocompromised individuals or persons with implanted medical devices such as catheters or voice prostheses ^56^ upon which the yeast grows as a biofilm ^7^.

Biofilms are structured communities of microorganisms attached to a surface. The cells are often encased within an extracellular matrix (ECM) which is commonly comprised of DNA ^8,9^, lipids ^8^, proteins ^8,10^ and polysaccharides ^8^. *C. albicans* is able to form biofilms on both abiotic and biotic surfaces and biofilm-associated cells are considerably more resistant to traditional antifungals when compared to planktonic cells ^11^. The reasons for this increased resistance are complex but include; the presence of an ECM which can act as a barrier to prevent antimicrobial agents reaching the cells ^12,13^, the presence of metabolically dormant persister cells inherent to biofilms ^14^, and the upregulation of drug efflux pumps ^15^. A significant percentage of human microbial infections arise from or are mediated via the formation of a biofilm ^16,17,18^, and this, combined the limited treatment options available, means the ability of *C. albicans* to grow as a biofilm is of particular medical interest.

*C. albicans* is a dimorphic fungus, it has the ability to undergo a morphogenic switch from a yeast, to pseudohyphal or hyphal forms in response to environmental cues. The virulence of *C. albicans* is closely linked with the capacity to switch between these forms; hyphal *C. albicans* cells are frequently located at sites of tissue invasion, and cells which are unable to readily form hyphae exhibit reduced virulence ^1^. The yeast-to-hyphal switch is also critical to biofilm formation as hyphal cells express a number of specific cell surface adhesins that enable cell-cell and cell-surface attachment ^19^. These adhesins, such as the agglutinin-like sequence (Als) proteins, possess a folded N-terminal domain required for protein-ligand interaction and a C-terminal peptide which covalently bonds to glycosylphosphatidylinositol (GPI) to anchor the adhesin in the fungal cell wall ^20^. The Als proteins also contain an amyloid forming region (AFR) in the N-terminal domain ^21^ which interacts with AFRs of other Als proteins. This results in the formation of large molecular weight clusters of Als proteins on the fungal cell wall called nanodomains which can bind multivalent ligands with high avidity ^22^. These nanodomains form in response to sheer forces applied to the adhesin molecules which cause the AFR to unfold and facilitate Als molecule aggregation ^23,24^. Nanodomains therefore strengthen adhesion and support the structure of mature biofilms ^25^.

*C. albicans* biofilm formation is a complex process involving tightly regulated, interwoven signalling pathways centrally controlled by a set of nine transcription factors; Bcr1, Brg1, Efg1, Flo8, Gal4, Ndt80, Rob1, Rfx2 and Tec1. These nine essential regulators function at different stages throughout *C. albicans* biofilm formation ^26^ and coordinate the expression of over 1000 target genes upregulated during biofilm formation ^27^. *C. albicans* biofilm formation can be divided into distinct stages that are governed by programmes of gene expression; attachment, initiation, maturation and dispersion. The attachment stage involves the initial attachment of *C. albicans* cells, primarily in the yeast-form ^28^, to a surface ^29^. Both nonspecific factors, such as cell surface hydrophobicity and electrostatic forces, and specific factors, such as adhesins on the yeast cell surface binding to precise ligands on the substratum to be colonised, are responsible for the preliminary attachment ^29^. Approximately 3-6 hours after the initial attachment, pseudohyphal and hyphal cells start forming from the proliferating yeast-form cells ^3^. This initiation step is characterised by the appearance of extracellular material. The maturation phase of biofilm growth lasts between 24-48 hours ^29^. Colonies of *C. albicans* continue to grow and secrete ECM, increasing the amount of material encasing the biofilm ^28^. The final stage of biofilm development is the dispersal stage during which yeast-form cells bud off from hyphal cells within a mature biofilm and disperse in order to establish additional biofilms elsewhere ^28,30^. The yeast-form cells emerging from mature biofilms have distinct characteristics compared to typical planktonic yeast cells; with enhanced adherence, an increased propensity to filament, and increased biofilm forming capability ^31^.

Elevated CO_2_ levels, as found in a number of physiologically relevant scenarios such as in exhaled breath or hypercapnia, have been shown to promote the yeast-to-hyphal switch in *C. albicans*. CO_2_ is converted to bicarbonate ions HCO_3_^-^ by the enzyme carbonic anhydrase which in turn activate the adenylate cyclase Cyr1, resulting in increased cAMP levels and the PKA dependent activation of hyphal specific genes ^32^. The yeast-to-hyphal switch is critical to the biofilm maturation process of *C. albicans* ^33^, as well as being important to its virulence ^1^. The effect of CO_2_ may be particularly important within the context of biofilm development on voice prostheses (VPs) since these devices are situated in the throat of patients where they are consistently exposed to high CO_2_ (5%) levels during exhalation. If CO_2_ does play a role in *C. albicans* biofilm maturation it could offer a possible explanation as to why *C. albicans* is found in such high frequencies on failed VPs. In addition, CO_2_ content within the blood is also elevated (46mmHg and 40mmHg for venous and arterial blood respectively versus 0.3mmHg found in atmospheric air) ^34,35^, and it has been estimated that as many as 80% of all microbial infections directly or indirectly involve pathogenic biofilms ^36^. Thus, the work presented here could be more widely applicable to bloodstream infections and biofilm formation within the body.

Here we demonstrate that physiologically relevant increases in CO_2_ accelerate *C. albicans* biofilm formation on silicone surfaces. Transcriptome analysis reveals that several core biofilm regulatory pathways, including those governed by Efg1, Bcr1, Brg1, and Ndt80, are upregulated. We also demonstrate that a high CO_2_ environment results in increased resistance of biofilms to azole antifungals, enhanced dispersal of cells from mature biofilms and an increase in capacity for glucose uptake. Moreover, a transcription factor knockout (TFKO) library screen demonstrated transcription factors involved in the acquisition of iron, such as the HAP transcription factors Hap43, Hap2, Hap3 and Hap5, to be important for *C. albicans* biofilm formation on silicone surfaces in atmospheric CO_2_ conditions. However, high CO_2_ was able to overcome the requirement for HAP transcription factor activity and enable *C. albicans* biofilms to forage for essential metabolites to support growth. Overall, we propose that *C. albicans* has adapted to utilise the high CO_2_ environment found in the host to promote its ability to colonise and to compete for nutrition. Our analysis reveals new approaches that can be taken to prevent *C. albicans* biofilm formation in high CO_2_ environments that pave the way for new therapeutic approaches to treat these highly drug resistant structures.

## Materials and Methods

### *Candida* strains and growth media

*Candida* strains (Supplementary Table S1) were routinely grown at 30°C in yeast peptone dextrose (YPD) media (2% peptone (BD Bacto), 2% D-glucose (Fisher Scientific), 1% yeast extract (BD Bacto)). For biofilm growth assays, *Candida* biofilms were grown at 37°C in RPMI-1640 media (Sigma-Aldrich, R8755) supplemented with 80*µ*g/ml uridine (Sigma-Aldrich, U3750) if required.

### *In vitro* biofilm growth assays

*C. albicans* biofilms were grown on a PDMS silicone elastomer (Provincial Rubber, S1). The silicone was cut into 1cm^2^ squares and placed in clips in a modified 24-well plate lid (Academic Centre for Dentistry Amsterdam, AAA-model) so they could be suspended in media within a 24-well plate. Silicone squares were incubated in 1ml 50% Donor Bovine Serum (DBS) (Gibco, 16030074) for 30 minutes at 30°C, then washed twice with 1ml PBS to remove excess DBS. 1ml of *C. albicans* were added to each well of a 24 well plate following resuspension in PBS at an OD_600_ of 1.0 and the lid with the silicone squares attached was placed on top so the silicone squares protrude into the cell suspension. Plates were then incubated at 37°C (in either 0.03% CO_2_ or 5% CO_2_) without shaking for 90 min to allow cell attachment to the silicone. After the attachment phase, the silicone squares were washed twice with 1ml PBS to remove any unattached cells and transferred to 1ml RPMI-1640 media (Sigma-Aldrich, R8755). They were then incubated at 37°C (in either 0.03% CO_2_ or 5% CO_2_) without shaking for 48h to allow biofilm maturation.

### Biofilm quantification via XTT assay

Biofilm growth was quantified using an XTT assay ^37^. Biofilms were washed twice with 1ml PBS to remove any planktonic cells before proceeding to quantification. After washing, the biofilms were transferred to a new pre-sterilised 24-well plate (Greiner Bio-one, CELLSTAR, 662160) containing 30*µ*g/ml XTT labelling reagent (Roche, 11465015001) and incubated at 37°C for 4h. After incubation, the biofilms were removed from the 24-well plate and the absorbance of the remaining XTT labelling reagent was measured at 492nm using a BMG LABTECH FLUOstar Omega plate reader machine.

### *C. albicans* transcription factor knockout (TFKO) screen

Biofilms using mutants from a *C. albicans* TFKO library ^38^ were seeded and grown for 48h on a PDMS silicone elastomer (Provincial Rubber, S1) as described above. Biofilm growth was quantified using the XTT assay. Experiments were performed in biological triplicate.

### Iron starvation of *C. albicans* biofilms

Biofilms were set up as described previously except they were incubated at 37°C in either 0.03% CO_2_ or 5% CO_2_ for 48h in RPMI-1640 containing varying concentrations of the Fe^2+^ chelator 3-(2-Pyridyl)-5,6-diphenyl-1,2,4-triazine-p,p’-disulfonic acid monosodium salt hydrate (Ferrozine – Sigma-Aldrich, 160601) or the Fe^3+^ chelator Deferasirox (Cambridge Bioscience, CAY16753-5). Ferrozine was made as a 100mM stock solution in sterile MQ H_2_O and diluted in RPMI-1640 to final concentrations of 250-500μM. Deferasirox was made as a 20mg/ml stock solution in DMSO and diluted in RPMI-1640 to final concentrations of 70-210*µ*g/ml. Relevant solvent controls were included. Final biofilms were quantified using an XTT assay.

### Preparation of PDMS-coated microscope slides

To prepare PDMS for coating microscope slides 16g (6.16 x 10^−4^ mol) silanol-terminated PDMS (cSt 1000, M_W_ 26000, from Fluorochem Ltd.) and 0.26g (12.48 x 10^−4^ mol, 1:4 stoichiometric ratio) cross-linking agent tetraethyl orthosilicate (TEOS – Sigma-Aldrich, 131903) were mixed at 3500rpm for 1 min using a DAC 150FV2-K speedmixer. At this point, 720*µ*l tin(II) ethylhexanoate (Sigma-Aldrich, S3252) made up at a concentration of 0.6M in toluene was added as a catalyst and the mix spun for a further 60 secs at 3500rpm. The elastomer mixture was then doctor bladed onto microscope slides using an automatic precision film applicator MTCX4 (Mtv-Messtechnik – blade width = 70mm, thickness adjustability 0-3000*µ*m). The doctor blade height was set 10*µ*m higher than the thickness of the microscope slide. The elastomer mix was poured over the top of the slide (with a bias towards the side of the microscope slide closest to the doctor blade), and then the doctor blade is moved at a constant speed over the substrate. The microscope slide was then air cured for 2h before being heat cured for 18h in a 70°C oven.

### Preparation of *C. albicans* biofilms on silicone coated slides for microscopy

Biofilms were grown directly on microscope slides that had been pre-coated with a PDMS silicone polymer (see above) for confocal imaging. Biofilms were grown with a prefabricated well. PDMS-coated microscope slides were incubated with 400*µ*l 50% Donor Bovine Serum (DBS) (Gibco, 16030074) in the wells for 30 min at 30°C and washed twice with 400*µ*l PBS. *C. albicans* overnight cultures were grown in YPD at 30°C and washed in PBS as described previously. 400*µ*l of the OD_600_ 1.0 standard cell suspension was added to the wells and incubated at 37°C (in either 0.03% CO_2_ or 5% CO_2_) without shaking for 90 min to allow cell attachment to the silicone surface of the microscope slide. After the attachment phase, the microscope slides were washed twice with 400*µ*l PBS to remove any unattached cells and then incubated at 37°C with 400*µ*l RPMI-1640 media in the wells for 6h, 24h or 48h (in either 0.03% CO_2_ or 5% CO_2_). Biofilms were washed twice with 400*µ*l PBS and then incubated in the dark for 45 min at 30°C in 400*µ*l PBS containing 50*µ*g/ml ConA-FITC (Sigma-Aldrich, C7642) and 20*µ*M FUN-1 (Invitrogen Molecular Probes, F7030). After incubation with the dyes, the stained biofilms were washed again with 400*µ*l PBS to remove any residual dye. The well was removed and 2 drops of ProLong™ Diamond Antifade Mountant (Invitrogen, P36965) was added to each stained biofilm. A cover slip was placed on top and the microscope slides were incubated in the dark at room temperature overnight to allow the mountant to cure.

### Confocal scanning laser microscopy (CSLM) of *Candida albicans* biofilms on silicone slides

Stained biofilms grown on silicone coated slides were imaged using a Zeiss LSM880/Elyra/Axio Observer.Z1 Confocal Microscope (Carl Zeiss Inc.) using the 488nm argon and the 561nm DP55 lasers. Images of the green (ConA-FITC) and the red (FUN-1) were taken simultaneously using a multitrack mode. Z-stacks were taken using the inbuilt ‘optimal’ settings to determine the optimal intervals (typically 1.5-2.0*µ*m slices) based upon sample thickness and magnification. The 20x and oil-immersion 40x objective lenses were used throughout. The image acquisition software used was ZENBlack and the image processing software was ZENBlue.

### RNA isolation from *C. albicans* biofilms

Total RNA was extracted in biological triplicate per condition (0.03% and 5% CO_2_) using the E.Z.N.A.™ Yeast RNA Kit (Omega Bio-Tek, R6870-01) as per the manufacturer’s instructions with a few modifications. Specifically, *C. albicans* CAI-4 biofilms were seeded and grown in 0.03% and 5% CO_2_ as described for *in vitro* biofilm growth assays. Mature biofilms were washed twice with 1ml ice-cold PBS to remove any planktonic cells. Biofilm cells were harvested by transferring silicone squares upon which the biofilms were growing into 5ml cold SE buffer/2-mercaptoethanol (provided in the E.Z.N.A.™ Yeast RNA Kit) and vortexing at 2500rpm for 1 min. The resulting biofilm cell suspension was pelleted by centrifugation at 4000rpm for 10 min at 4°C. The supernatant was discarded and the cells re-suspended in fresh 1ml cold SE buffer/2-mercaptoethanol, this cell suspension was transferred to a 2ml Eppendorf tube. The cell suspension was centrifuged again for 10 min at 4°C, the supernatant discarded and the pellet re-suspended in 480*µ*l fresh SE buffer/2-mercaptoethanol. The biofilm cell suspension was incubated with 80*µ*l lyticase stock solution (5000units/ml in SE buffer) at 30°C for 90 min. The resulting spheroplasts were pelleted by centrifugation at 2900rpm for 10 min at 4°C and the supernatant aspirated and discarded. The spheroplasts were gently re-suspended in 350*µ*l YRL buffer/2-mercaptoethanol (provided in the E.Z.N.A.™ Yeast RNA Kit). The rest of the RNA extraction proceeded as per the manufacturer’s instructions including the optional DNase digestion step. RNA was eluted in 30*µ*l DEPC water, the concentration and purity established using a NanoDrop ND-1000 spectrophotometer (NanoDrop Technologies) and stored at - 80°C.

### Library Preparation and RNA Sequencing

RNA samples were sent to the Centre for Genome Enabled Biology and Medicine (Aberdeen, UK) for library preparation and sequencing. Before library preparation, the quality and quantification of RNA samples were evaluated with TapeStation (Agilent) and Qubit (Thermal Fisher). Samples with a minimum RIN of 8.0 proceeded to library preparation. The input of RNA was based on the specifically measured RNA concentration by Qubit. The mRNA-Seq libraries were prepared using TruSeq™ Stranded mRNA Sample Preparation Kit (Illumina) according to the manufacturer’s instructions. Briefly, Poly-A RNA were purified from 500ng of total RNA with 1ul (1:100) ERCC spike (Thermal Fisher) as an internal control using RNA purification oligo(dT) beads, fragmented and retrotranscribed using random primers. Complementary-DNAs were end-repaired, and 3-adenylated, indexed adapters were then ligated. 15 cycles of PCR amplification were performed, and the PCR products were cleaned up with AMPure beads (Beckman Coulter). Libraries were validated for quality on Agilent DNA1000 Kit and quantified with the qPCR NGS Library Quantification kit (Roche). The final libraries were equimolar pooled and sequenced using the High Output 1×75 kit on the Illumina NextSeq500 platform producing 75bp single-end reads. For each library a depth of 50-70M reads was generated.

### Analysis of RNA-Seq data

Analysis of RNA-Seq data was performed using the Galaxy web platform ^39^. The quality of the RNA sequencing reads was checked using FastQC v0.11.5 ^40^ with default settings. Low quality ends (Phred score < 20) and any adaptor sequences were trimmed using TrimGalore! v0.4.3 ^41^. Reads which became shorter than 40bp after trimming were removed from further analysis. After trimming, 97.7% of initial reads remained and the quality was checked again using FastQC v0.11.5 ^40^. There were no Poly-A reads (more than 90% of the bases equal A), ambiguous reads (containing N) or low quality reads (more than 50% of the bases with a Phred score < 25). After processing, the mean Phred score per read was 35. Processed reads were aligned with the reference *C. albicans* genome SC5314 version A21-s02-m09-r10 using HISAT2 v2.1.0 ^42^ with single-end reads and reverse strand settings (rest of the settings were default). After alignment, the number of mapped reads which overlapped CDS features in the genome (using the *C. albicans_SC5314_version_A21-s02-m09-r10_features.gtf* annotation file ^43^) were determined using htseq-count v0.9.1 ^44^ with default settings. Reads aligning to multiple positions or overlapping more than one gene were discarded, counting only reads mapping unambiguously to a single gene. Differential gene expression analysis between conditions (0.03% and 5% CO_2_) was performed using DESeq2 v1.18.1 ^45^ with default settings.

### Gene Set Enrichment Analysis of transcription profiles

Downstream analysis of RNA Sequencing data was performed using the PreRanked tool of Gene Set Enrichment Analysis (GSEA; Broad Institute) ^46^ which compares a pre-ranked significantly differentially expressed gene list to a functional gene set database. False discovery rate (FDR) q-values were calculated based upon 1000 permutations. The gene set database used was assembled by Sellam, A. *et al*. as described in ^47^, which is based upon experimental analyses from published studies, Gene Ontology (GO) term categories curated by the *Candida* Genome Database ^48^ and protein-protein interaction information derived from *Saccharomyces cerevisiae* data curated by the *Saccharomyces* Genome Database ^49^. Gene set networks were generated in Cytoscape 3.7.1 (Available at: https://cytoscape.org/) ^50^ using the EnrichmentMap plug-in (Available at: http://apps.cytoscape.org/apps/enrichmentmap). Gene expression heat maps based on GO term categories were created using the Pheatmap package in R Studio.

### Antifungal treatment of *C. albicans* biofilms

Biofilms were set up as described previously and grown in RPMI-1640 for 24h at 37°C. The biofilms were then transferred to fresh RPMI-1640 media containing a select antifungal. Three antifungals were tested; Fluconazole, Miconazole and Nystatin. Fluconazole (Santa Cruz Biotechnology, sc-205698) was made as a 50mg/ml stock solution in ethanol and diluted in RPMI-1640 final concentrations ranging from 8-256*µ*g/ml. Miconazole (Santa Cruz Biotechnology, sc-205753) was made as a 50mg/ml stock solution in DMSO and also diluted in RPMI-1640 to final concentrations ranging from 8-256*µ*g/ml. Nystatin (Santa Cruz Biotechnology, sc-212431) was made as a 5mg/ml stock solution in DMSO and diluted in RPMI-1640 to final concentrations ranging from 1-8*µ*g/ml. Drug vehicle controls (0.5% ethanol for Fluconazole, 0.5% DMSO for Miconazole and Nystatin) were used in all cases. The biofilms matured in the RPMI-1640 media containing the select antifungal for a further 24h at 37°C in both 0.03% and 5% CO_2_ before proceeding to quantification via the XTT assay. Experiments were performed in biological and technical triplicate.

### *C. albicans* attachment assay

CAI-4 cells were seeded onto silicone-coated microscope slide for 90 min as described previously for growing biofilms for confocal microscopy, except here an OD_600_ of 0.1 was used instead of 1.0 and the silicone-coated microscope slides were not pre-incubated with DBS. The slide surface was washed twice with 400*µ*l PBS to remove any unattached cells. Images were taken using a Leica DMR fitted with a Leica DFC9000 GT camera using a 20x objective lens and brightfield settings. The image acquisition software used was the Leica Application Suite X package. Using identical microscope settings throughout, five images were taken of each of three biological replicates in both 0.03% and 5% CO_2_ and the cells per image counted using the ‘Cell Counter’ function in ImageJ.

### 2-deoxyglucose (2-DG) treatment of *C. albicans* biofilms

Biofilms were set up as described previously except they were incubated at 37°C in either 0.03% CO_2_ or 5% CO_2_ for 48h in RPMI-1640 containing varying concentrations of 2-DG. 2-DG (Sigma-Aldrich, D6134) was made up as a 20% stock solution in sterile MQ H_2_O and diluted in RPMI-1640 to final concentrations of 0.25-1%. Biofilms were quantified using XTT assays and images were also taken. Experiments were performed in biological and technical triplicate.

### *C. albicans* biofilm dispersion assay

Biofilms were set up as described previously and grown in RPMI-1640 for 48h at 37°C. The spent media, containing dispersed cells, was sonicated at amplitude 4*µ*m for 10 secs to separate clumps of hyphal cells (previous work in our lab has demonstrated these sonication settings do not affect viability). After sonication, the dispersed cells were diluted 1 in 10 and 200*µ*l of this suspension was plated on YPD agar plates in triplicate. YPD plates were incubated for 48h at 37°C to allow colonies to form, at which point the number of colonies were manually counted using a Stuart Scientific Colony Counter. Experiments were performed in biological and technical triplicate.

## Results

### CO_2_ enhances *C. albicans* biofilm formation

*C. albicans* biofilms were seeded and grown on silicone in CO_2_ levels found in exhaled air (5%) and atmospheric air (0.03%) for 24 and 48 hours. Biofilms were quantified using the XTT assay which acts as a readout of cell number ^37^. After 24 hours of growth, both CAI4pSM2 and SN250 *C. albicans* strains exhibited a significantly higher cell number within biofilms grown in a 5% CO_2_. However, after 48 hours, cell numbers within biofilms grown in both CO_2_ conditions appeared equal (Figure 1A). Interestingly, although cell number appeared equivalent at 48h, the resultant biofilm mass appeared noticeably larger in biofilms grown under elevated CO_2_ conditions (Figure 1B).

**Figure 1:**
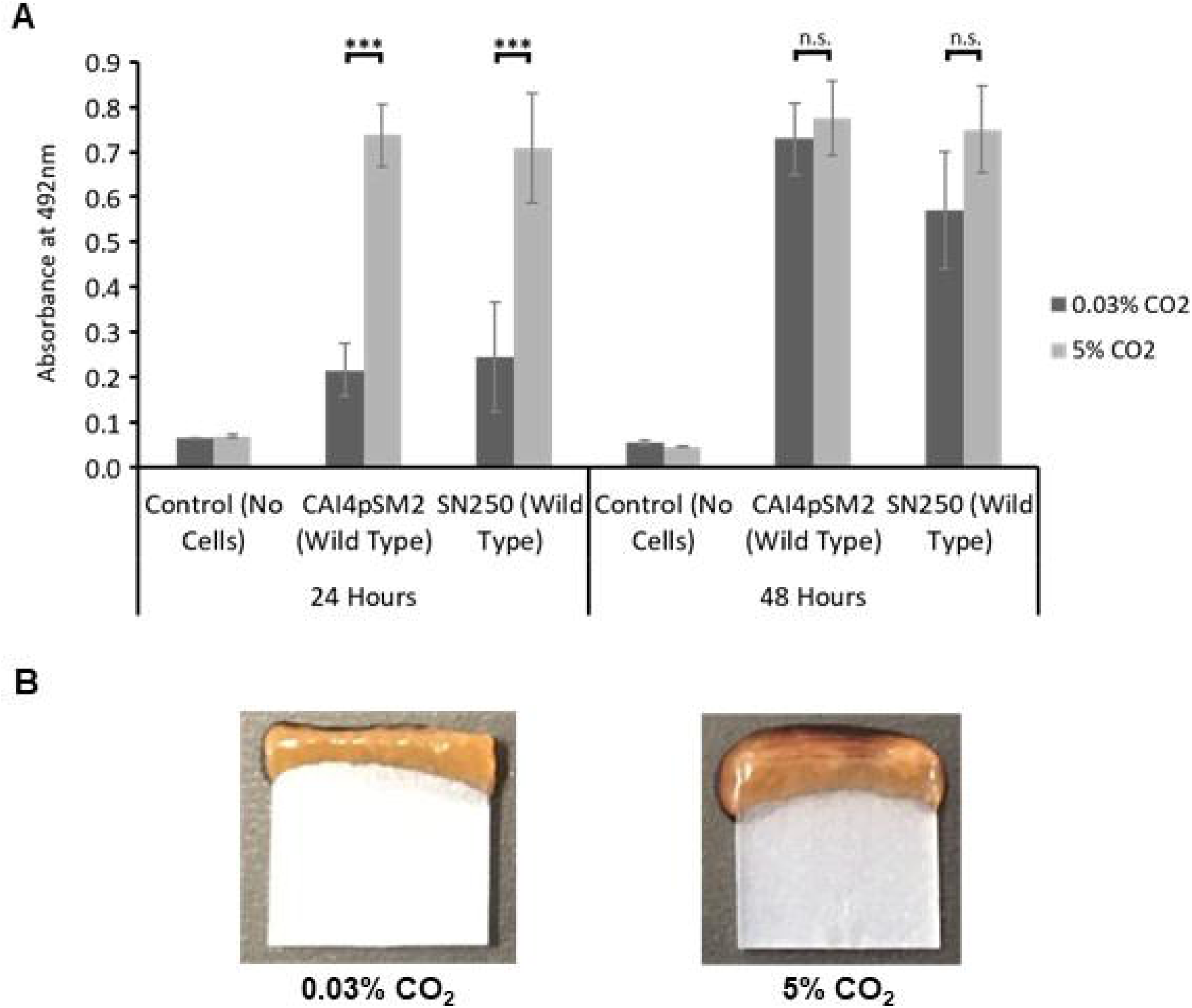
The effect of high CO_2_ (5%) on *C. albicans* biofilm formation. **(A)** Biofilms were seeded and grown for 24h or 48h in 0.03% or 5% CO_2_, the resulting biofilms were quantified using the XTT assay with absorbance at 492nm as a readout. The graph represents three biological replicates each containing technical triplicates, error bars denote Standard Deviation. Paired two-tail t-tests were carried out: *p<0.05, **p<0.01, ***p<0.001, n.s. = not significant. **(B)** Representative images of *C. albicans* (SN250 strain) biofilms grown in 0.03% and 5% CO_2_ for 48h.

### Analysis of the effects of CO_2_ on phases of *C. albicans* biofilm formation

We sought to determine which phases of biofilm growth were influenced by CO_2_ elevation. To investigate the attachment phase *C. albicans* cells were seeded for 90 min onto silicone-coated microscope slides under 0.03% CO_2_ or 5% CO_2_ levels. We observed that exposure to elevated CO_2_ led to a significant increase in the number of cells that attached to silicone (mean of 2108 cells in 0.03% vs. a mean of 7033 cells in 5% CO_2_) (Figure 2A). Cells also appeared to attach as larger aggregations in 5% CO_2_ when compared to those in 0.03% CO_2_ (Supplementary Figures S1A and B) indicating that both cell-substrate and cell-cell attachments may be enhanced.

**Figure 2:**
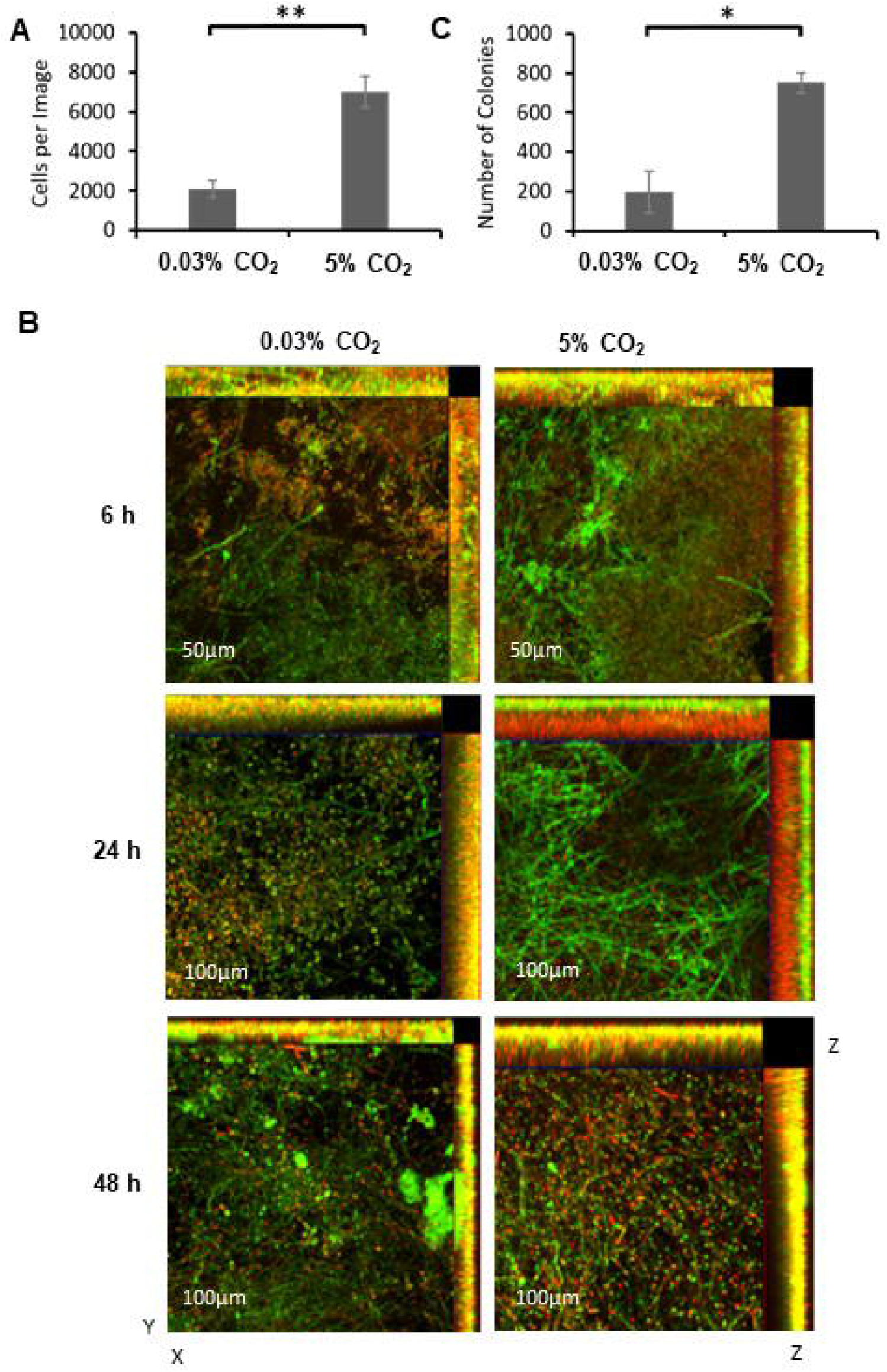
The effects of CO_2_ on *C. albicans* biofilm growth. **(A)** <u>Attachment:</u> *C. albicans* CAI-4 cells were seeded onto silicone-coated microscope slides under 0.03% or 5% CO_2_ and images taken at 20x objective magnification. Cells per image were counted and the mean calculated across three biological replicates (five images per replicate). A paired two-tail t-test carried out: **p<0.01. Error bars denote Standard Deviation. **(B)** <u>Maturation:</u> Biofilms were seeded on silicone-coated microscope slide and grown for 6h, 24h and 48h. Biofilms were stained with ConA-FITC (green) and FUN-1 (red). Z-stack Images were taken using 20x (6h) and 40x (24 and 48h) magnifications. Experiments were repeated in triplicate and representative maximum intensity images are presented as well as a z stack profiles. **(C)** <u>Dispersion:</u> Spent media was collected from biofilms grown for 48h in 0.03% or 5% CO_2_ and diluted 1:10 before being plated to assess the number of colonies. Three biological replicates each containing technical triplicates were conducted, error bars denote Standard Deviation. A paired two-tail t-test was carried out: *p<0.05.

Confocal scanning laser microscopy (CSLM) was used to investigate the effects of CO_2_ on biofilm growth during maturation. *C. albicans* biofilms were seeded on silicone-coated microscope slides and images were taken at 6h, 24h and 48h of growth under either 0.03% or 5% CO_2_ growth conditions (Figure 2B). Biofilm images are displayed as maximum intensity ortho-projections of Z-stacks to give a view of the overall structures of the biofilms. After 6h growth in 0.03% CO_2_, the majority of cells were found in the yeast form with some visibly initiating hyphae. In comparison, biofilms grown in 5% CO_2_ appeared to consist of a high proportion of hyphal cells, were visibly denser and had begun to exhibit an ordered structure in the Z-plane (Figure 2B). After 24h growth in 0.03% CO_2_, biofilms were progressing through the maturation stage with the appearance numerous hyphal cells. However, the 5% CO_2_ biofilms displayed a fully mature biofilm organisation displaying hyphal cells organised in a brush-like structure above a basal layer of yeast cells (Figure 2B). At 48h, biofilms grown in both 0.03% and 5% CO_2_ appeared as dense mature structures, however, biofilms grown under elevated CO_2_ appeared larger (Figure 2B) as had been observed macroscopically (Figure 1B).

Dispersion is the final stage of biofilm formation, we therefore investigated whether CO_2_ elevation resulted in increased levels of cell shedding. We routinely observed that spent RPMI-1640 media isolated after biofilm growth in 5% CO_2_ contained more cells than that of taken from biofilms grown in 0.03% CO_2_ (Supplementary Figure S2A). We quantified this by seeding and growing *C. albicans* biofilms for 48h in 0.03% and 5% CO_2_ and conducting colony forming unit (CFU) assays using the spent RPMI-1640 media (Figure 2C and Supplementary Figure S2B). Our results showed an approximate four-fold increase in cell number released from mature biofilms when grown under elevated CO_2_ conditions, consistent with an increase in dispersal. Overall, these data demonstrate that the elevation of CO_2_ enhances each stage of the *C. albicans* biofilm forming process, from attachment through maturation to dispersion.

### Identification of the regulatory mechanisms that govern CO_2_ acceleration of *C. albicans* biofilm formation

In planktonic *C. albicans* cells CO_2_ is converted to bicarbonate ions (HCO_3_^-^) which stimulates the adenylate cyclase Cyr1 (Cdc35), an increase in cAMP and activation of PKA ^32^. We investigated whether CO_2_ elevation may drive biofilm formation and maturation via a similar Ras/cAMP/PKA mechanism. We conducted biofilm growth assays using *C. albicans* mutants lacking key components of the pathway; *ras1Δ/Δ, cdc35Δ/Δ, CDC35*^ΔRA^ (adenylate cyclase missing the Ras1 interacting domain), *tpk1Δ/Δ* (missing a catalytic subunit isoform of PKA), and *tpk2Δ/Δ* (missing the other catalytic subunit isoform of PKA). Biofilm formation was quantified after 48h of growth and compared to an isogenic wild type control. The *ras1Δ/Δ* mutant displayed significantly attenuated biofilm growth in 0.03% CO_2_ but this was rescued to wild type levels in 5% CO_2_ (Figure 3A), indicating that Ras1 function is dispensable for biofilm formation under conditions of elevated CO_2_. The *cdc35Δ/Δ* and *CDC35*^ΔRA^ mutants both exhibited significantly reduced biofilm growth when grown under either atmospheric or elevated CO_2_ conditions (Figure 3A). Conversely, both *tpk1Δ/Δ* and *tpk2Δ/Δ* mutants exhibited biofilm growth equivalent to the wild type under both CO_2_ conditions (Figure 3A). Taken together, this data implies the CO_2_-mediated effect on *C. albicans* biofilm growth is reliant on Cyr1 but can bypass a requirement for Ras1 and that the PKA isoforms Tpk1 and Tpk2 are functionally redundant with respect to cAMP activation of the biofilm programme. Our findings are consistent with the adenylate cyclase Cyr1 as a key CO_2_ sensor in the enhanced biofilm growth observed under elevated CO_2_ conditions.

**Figure 3:**
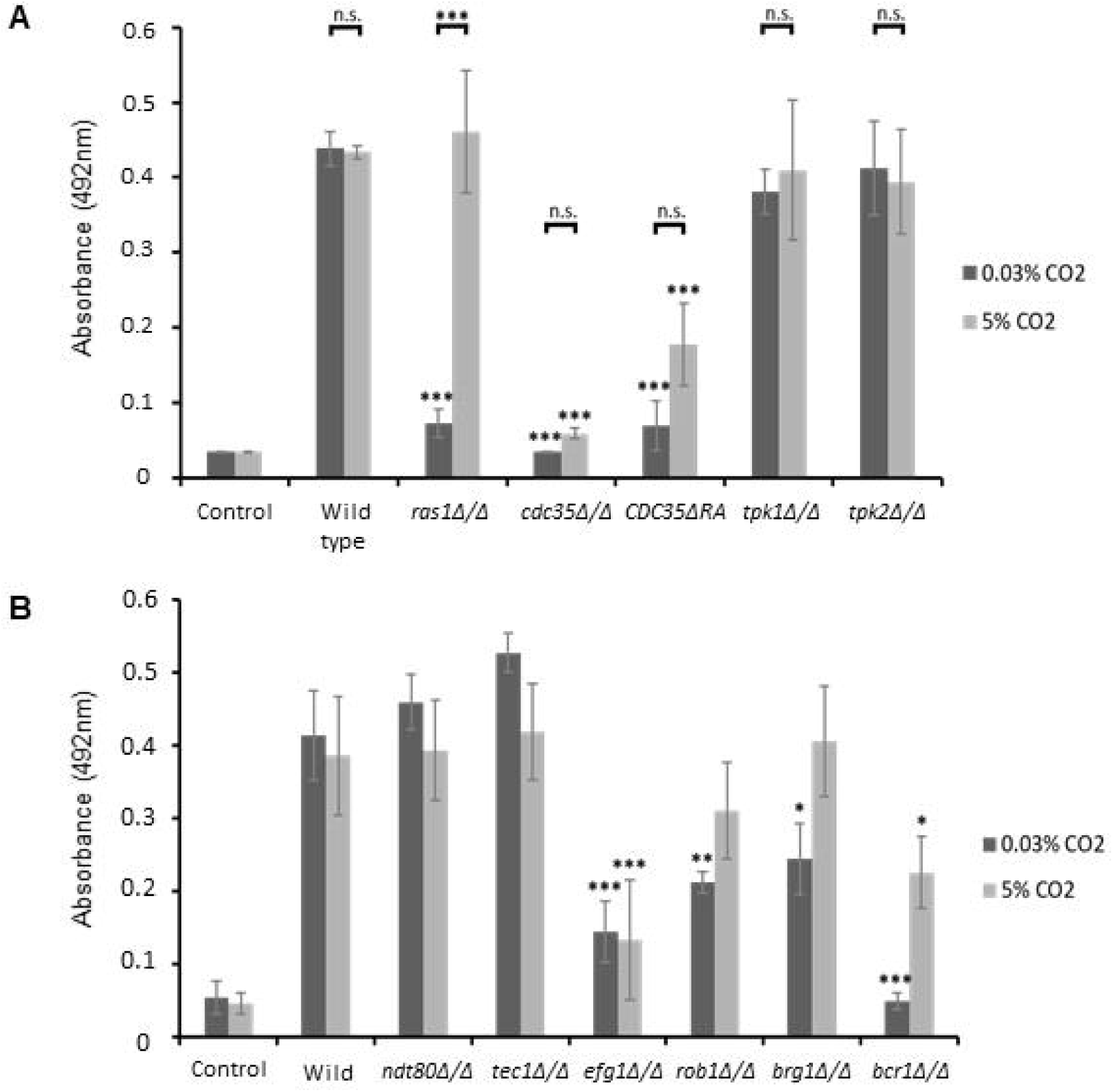
Biofilm growth assays of *C. albicans* Ras1-Cyr1-PKA pathway and central biofilm regulator null mutants. Cells were seeded and grown as biofilms for 48h before XTT quantification. Control wells with no cells were included. **(A)** Graph represents three biological replicates each containing technical triplicates, error bars denote Standard Deviation. **(B)** Graph represents three biological replicates, error bars denote Standard Deviation. Two-way ANOVAs followed by Tukey tests for multiple comparisons were carried out: *p<0.05, **p<0.01, ***p<0.001, n.s. = not significant. Stars directly above the bars indicate a significant difference to the wild type in the same CO_2_ environment.

To investigate the molecular basis of the activation of *C. albicans* biofilm formation in 5% CO_2_, we conducted a screen of an available transcription factor knockout (TFKO) library ^38^ containing 147 mutants each lacking a non-essential transcription factor. This screen identified 122 deletions that had no effect upon biofilm formation in either CO_2_ condition (Supplementary Figure S3) and 22 transcription factors which attenuated biofilm formation (Supplementary Table S2). Six TFKO mutants (*tup1Δ/Δ, sef1Δ/Δ, swi4Δ/Δ, pho4Δ/Δ, bcr1Δ/Δ* and *efg1Δ/Δ*) had diminished biofilm growth in both 0.03% and 5% CO_2_, 12 had reduced biofilm formation only in 0.03% CO_2_ (*hap2Δ/Δ, rbf1Δ/Δ, rob1Δ/Δ, fgr15Δ/Δ, dal81Δ/Δ, mig1Δ/Δ, brg1Δ/Δ, C4_00260WΔ/Δ, zcf27Δ/Δ, C1_13880CΔ/Δ, crz1Δ/Δ*, and *hap43Δ/Δ*), and 4 TFKOs had reduced biofilm formation only in 5% CO_2_ (*leu3Δ/Δ, mbp1Δ/Δ, bas1Δ/Δ*, and *try6Δ/Δ*) (Supplementary Table S2 and Supplementary Figure S3). Intriguingly, 3 additional mutants, *zcf17Δ/Δ, zcf30Δ/Δ* and *mac1Δ/Δ*, displayed increased biofilm growth in 0.03% CO_2_ but no significant difference in 5% CO_2_.

We performed gene ontology enrichment analysis on the genes encoding the 25 transcription factors whose loss impacted upon biofilm growth to group them according to biological processes. This revealed that 7 of these transcription factors – Brg1, Bcr1, Efg1, Rob1, Dal81, Leu3, and Try6 – were already known to be involved in the regulation of single-species biofilm formation within *C. albicans* (Table S2). Interestingly, out of these, only the *efg1Δ/Δ* and *bcr1Δ/Δ* mutant exhibited attenuated biofilm growth in both 0.03% and 5% CO_2_. The *brg1Δ/Δ, dal81Δ/Δ*, and *rob1Δ/Δ* mutants had significantly reduced biofilm growth in 0.03% CO_2_ but this was rescued in the 5% CO_2_ environment (Supplementary Table S2), possibly indicating redundancy of these TFs when cells are exposed to high CO_2_. This important finding suggests that CO_2_ elevation can bypass the requirement for some of the key transcriptional regulators of biofilm formation (Figure 3B). The *leu3Δ/Δ* and *try6Δ/Δ* mutants had significantly reduced biofilm growth in 5% CO_2_ but no significant difference in 0.03% CO_2_. Eight of the TFs identified – Brg1, Crz1, Efg1, Mac1, Mig1, Rob1, Zcf27, Zcf30 – are associated with the positive regulation of filamentous growth and Tup1 is involved in the negative regulation of filamentous growth. *C. albicans* biofilm formation is strongly linked with the yeast-to-hyphal switch, with mutants unable to perform this switch having previously been shown to be deficient in biofilm growth ^33^.

The TFKO screen also revealed that mutants lacking transcription factors associated with metal ion homeostasis, specifically iron homeostasis, had altered biofilm-formation in 0.03% and/or 5% CO_2_ (Figure 4A and Supplementary Table S2). Mutants lacking genes expressing components of the HAP complex; *hap2Δ/Δ, hap3Δ/Δ, hap5Δ/Δ*, and *hap43Δ/Δ*, exhibited significantly reduced biofilm growth after 48h compared to wild type in 0.03% CO_2_. However, their biofilm growth was significantly higher in 5% CO_2_, reaching wild type levels in the cases of the *hap3Δ/Δ, hap5Δ/Δ*, and *hap43Δ/Δ* mutants (Figure 4A). This is an important observation as it indicates that an increase in CO_2_ concentration is sufficient to compensate for the loss of these transcription factors. The HAP complex in *C. albicans* is responsible for the regulation of iron homeostasis ^51^. Sef1 acts downstream of this complex and the *sef1Δ/Δ* mutant displayed significantly reduced biofilm growth in both 0.03% and 5% CO_2_ after 48h growth (Figure 4A). These data suggest that the elevation of CO_2_ can bypass the requirement for HAP complex activity in biofilm formation in a Sef1 dependent manner.

**Figure 4:**
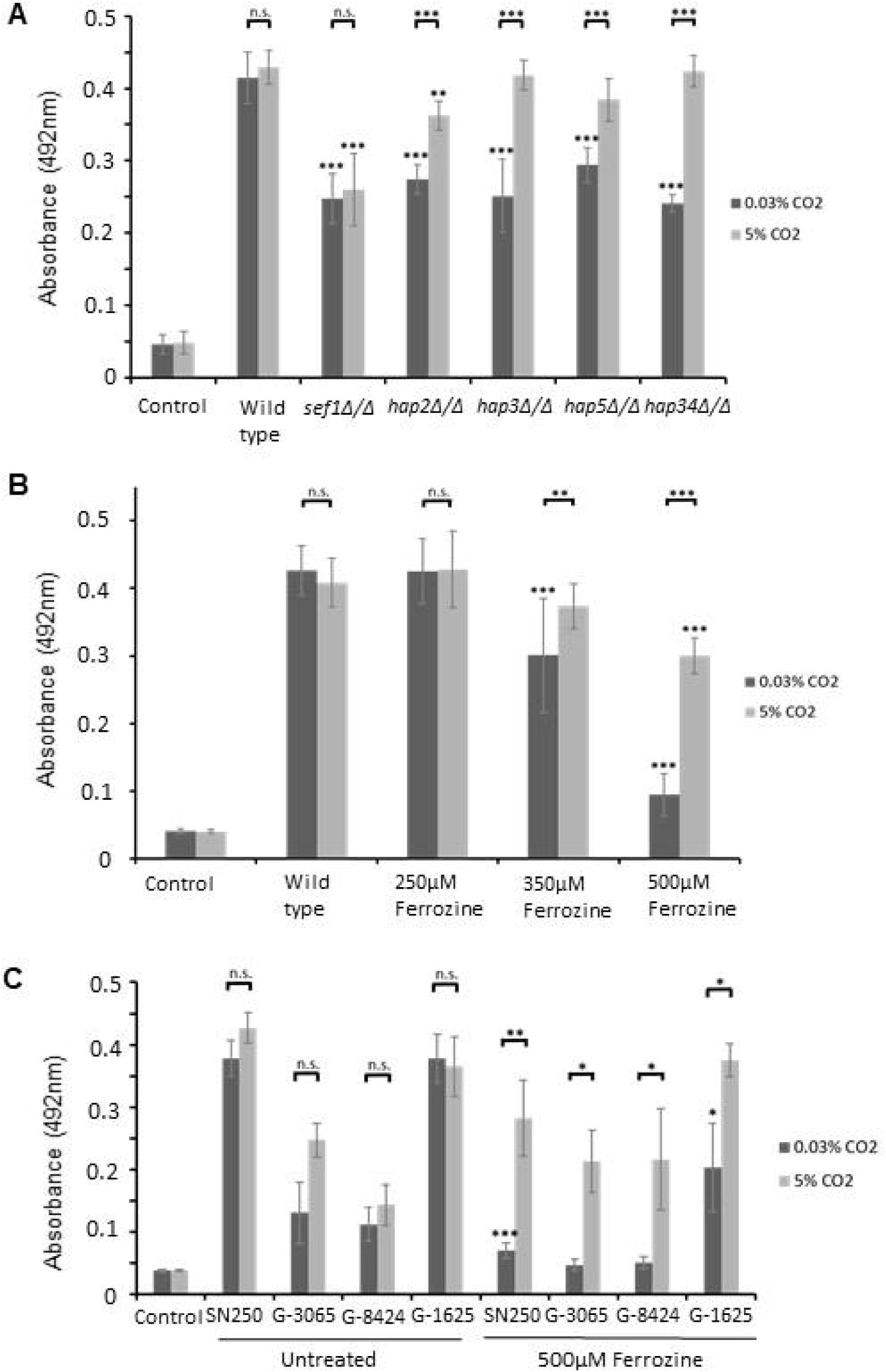
The effect of high (5%) CO_2_ on iron homeostasis in *C. albicans* biofilms. Biofilms were seeded and grown for 48h before XTT quantification. Control wells with no cells were set up as media controls. **(A)** Biofilm growth assay using TFKO mutants lacking iron homeostatic transcription factors. **(B)** Iron starvation biofilm growth assay of in the presence of the Fe^2+^ chelator Ferrozine. **(C)** Iron starvation biofilm growth assay of clinical isolates in the presence of Ferrozine. Graph represents three biological replicates each containing technical triplicates, error bars denote Standard Deviation. Two-way ANOVAs followed by Tukey tests for multiple comparisons were carried out: *p<0.05, **p<0.01, ***p<0.001, n.s. = not significant.

To further determine whether elevated CO_2_ enhanced *C. albicans* ability to appropriate iron from the environment biofilms were grown in the presence of the Fe^2+^ chelator 3-(2-Pyridyl)-5,6-diphenyl-1,2,4-triazine-p,p’-disulfonic acid monosodium salt hydrate (Ferrozine). Iron chelation was observed to have a marked effect on biofilm growth at and above 350μM Ferrozine (Figure 4B). As had been observed with TFKO strains of the HAP complex, the elevation of CO_2_ counteracted the effects of iron limitation (Figure 4B) providing further evidence of a role for CO_2_ in enhancing iron uptake or utilisation capability. This tolerance to iron starvation of *C. albicans* biofilms grown in 5% CO_2_ was also exhibited by *tpk1Δ/Δ* and *tpk2Δ/Δ* mutants (Supplementary Figure S4), indicating these PKA isoforms are functionally redundant with respect to iron homeostatic pathways. Clinical strains of *C. albicans* isolated from failed voice prostheses displayed the same phenotype with biofilm formation rescued in 5% CO_2_ when grown in the presence of 500μM Ferrozine (Figure 4C). A transcriptomic analysis comparing biofilms grown in 0.03% and 5% CO_2_ (described below) revealed that several genes related to iron acquisition were upregulated in 5% CO_2_ biofilms, providing an explanation for the tolerance to iron sequestration in high CO_2_ (Figure 4). For example, *FTR2*, which encodes the high affinity iron permease Ftr2, had increased expression. In addition, *CFL4* which encodes a putative ferric reductase, is regulated by Sef1 and induced in low-iron conditions ^52^ was also upregulated in high CO_2_ biofilms after 48h. Finally, the *CSA2* and *RBT5* genes, which encode proteins involved in the acquisition of iron from haem groups, were also upregulated. Csa2 and Rbt5 have both previously been found to be required for normal biofilm formation in RPMI-1640 media ^53,54^.

### Transcriptome analysis of *C. albicans* biofilms grown in high and low CO_2_

To further investigate the effect of high CO_2_ on *C. albicans* biofilm growth, we performed an RNA Sequencing analysis of *C. albicans* biofilms grown in 0.03% and 5% CO_2_. Growth in 5% CO_2_ led to a global response resulting in the significant differential expression (false-discovery-rate adjusted p-value (q) ≤ 0.05) of 2875 genes, with roughly equal numbers of genes up- and downregulated (1441 up and 1434 down) (Supplementary Figure S5A). 80 genes were strongly (log2 fold change >2) upregulated and 25 were strongly (log2 fold change <-2) downregulated. To investigate the cellular pathways which these differentially expressed genes are within we conducted Gene Set Enrichment Analysis (GSEA; Broad Institute) ^46^. GSEA revealed that biofilm formation pathways controlled by four of the nine ‘core’ biofilm regulator transcription factors ^27,55^ were upregulated in 5% CO_2_ biofilms at 48h (Figure 5A). Genes downstream of Brg1, Efg1, Ndt80 and Bcr1 are enriched in the upregulated genes at the top of the ranked list of differentially expressed genes (NES = +2.79, +2.66, +2.56, and +2.60 respectively) (Figure 5A), indicating high CO_2_ drives expression of genes previously described as important in the biofilm-forming ability of *C. albicans*.

**Figure 5:**
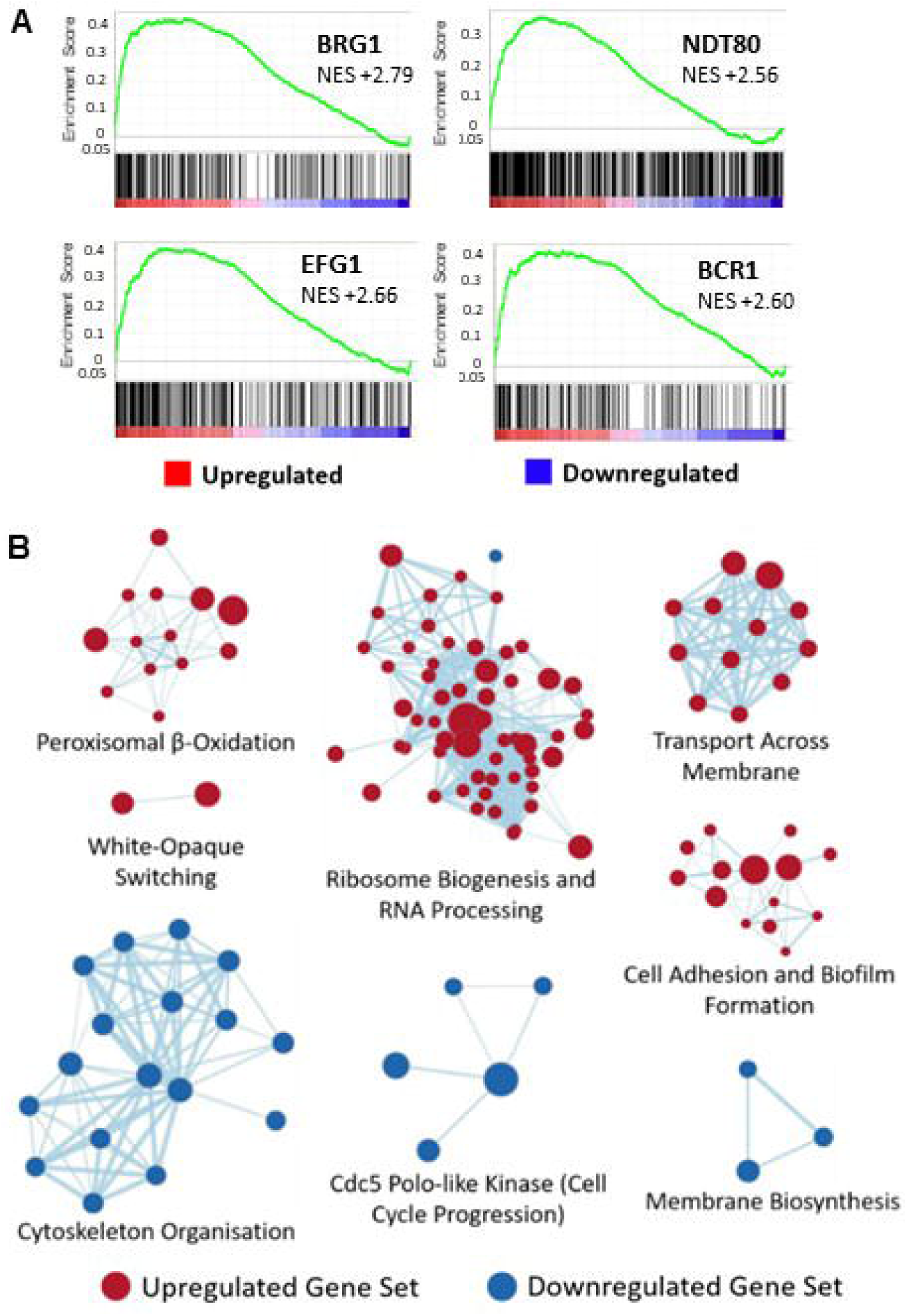
Global gene expression changes in 5% CO_2_ vs. 0.03% CO_2_ *C. albicans* biofilms. **(A)** GSEA enrichment plots of central biofilm regulator gene sets with altered expression levels as assessed by RNA Sequencing, four of the nine identified core regulators of biofilm formation (Brg1, Efg1, Ndt80, and Bcr1 ^27^) were identified as having positive GSEA scores. Vertical black lines represent individual genes in the significantly differentially expressed ranked gene list from upregulated (left) to downregulated (right). The enrichment score increases if there are lots of genes towards the beginning of the ranked list (upregulated). NES = normalised enrichment score, positive NES indicates enrichment in the upregulated group of genes. **(B)** Gene set cluster map showing the most upregulated and downregulated gene sets as determined by GSEA along with their cellular functions. Each circle is a gene set and the lines between them depict how much they overlap, thicker line = more genes in common.

As GSEA can show enrichment profiles exhibiting correlations with several overlapping gene sets, we visualised networks of similar gene sets using the Cytoscape 3.7.1 EnrichmentMap plug-in ^50^. Upregulated gene sets included those encoding membrane transporters, ribosome biogenesis, peroxisomal β-oxidation and white-opaque switching (Figure 5B). Pathways involved in cytoskeleton organisation, cell-cycle progression and membrane biosynthesis were downregulated in high CO_2_ conditions (Figure 5B), indicating cells within biofilms in high CO_2_ may have stopped dividing at the 48h time point assessed, consistent with such biofilms having moved more rapidly through the maturation process.

GSEA suggested that cell adhesion processes were upregulated in high CO_2_ biofilms (NES = +2.13) (Figures 6A). GO Slim process analysis also highlighted many genes associated with cell adhesion were upregulated in *C. albicans* biofilms grown in 5% CO_2_ after 48h (Figure 6B). The second highest upregulated gene in 5% CO_2_ biofilms compared to 0.03% CO_2_ biofilms was *ALS1* with a 3.77 log2 fold change (Figure 6B). The Als1 cell surface adhesin has previously been shown to have important roles in biofilm formation ^56^, and its expression is controlled by the biofilm transcription regulation network composed of Brg1, Rob1, Tec1, Ndt80, Bcr1 and Efg1 ^27^. Other genes encoding cell surface adhesins such as *ALS4* and *ALS2* were also upregulated in 5% CO_2_ biofilms after 48h (Figure 6B).

**Figure 6:**
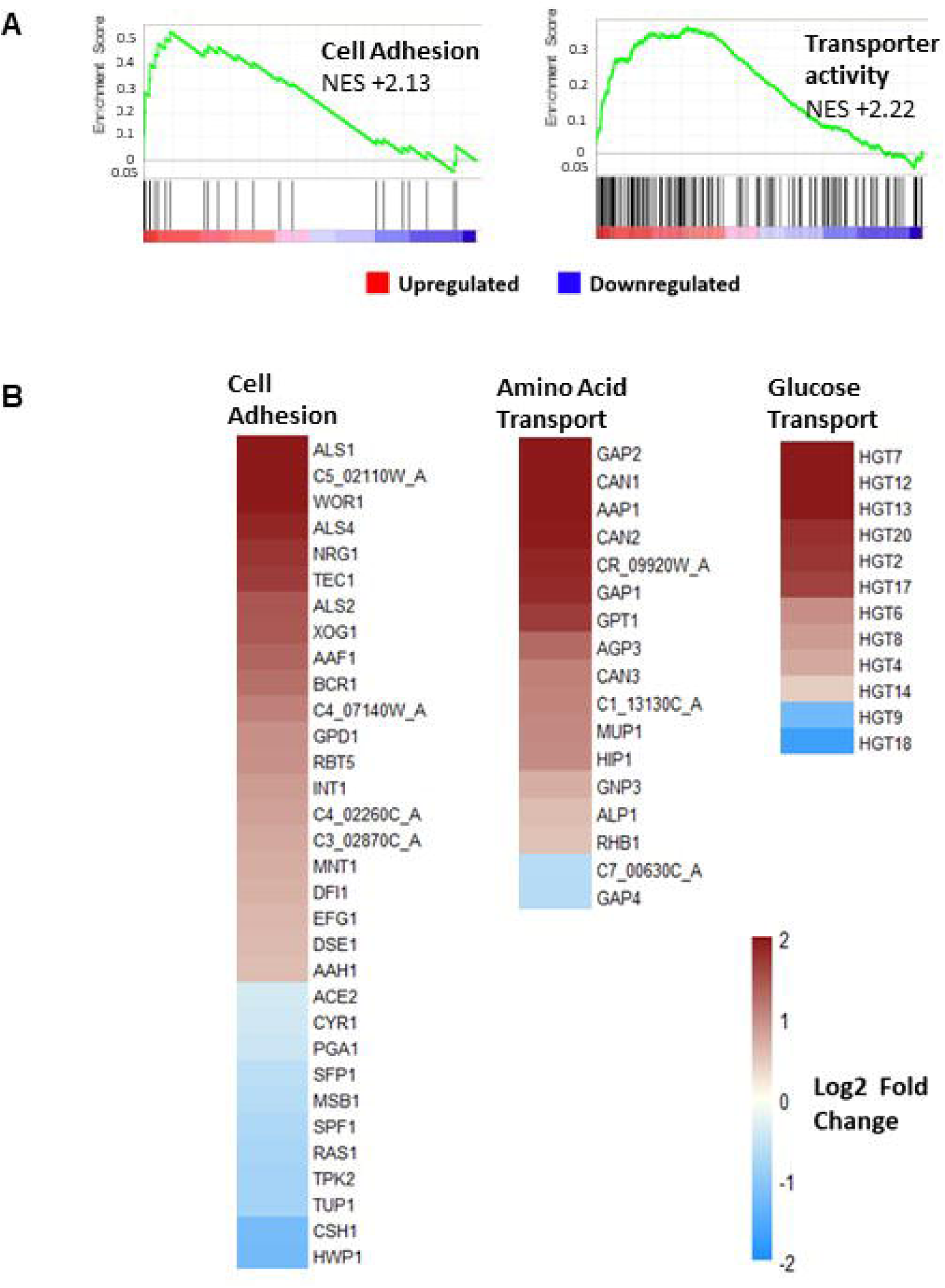
Adhesion and transport processes are upregulated in 5% CO_2_ *C. albicans* biofilms. **(A)** GSEA enrichment plot of the BIOLOGICAL ADHESION_BIO and TRANSPORTER ACTIVITY_MOL, AMINO ACID TRANSPORT_BIO and CARBOHYDRATE TRANSPORTER ACTIVITY_MOL gene sets containing genes under the GO terms ‘transporter activity’ and ‘amino acid transport’ NES = normalised enrichment score, positive NES indicates enrichment in the upregulated group of genes. **(B)** Heat map of significantly differentially expressed genes associated with cell adhesion, amino acid transport and glucose transport as identified by GO Slim process analysis. The colours saturate at log2 fold change of 2 and -2.

GSEA has also identified that genes involved in membrane transporter activity are enriched in the upregulated genes at the top of the ranked list of differentially expressed genes (NES = +2.22) (Figure 6A). Specifically, genes encoding amino acid transporters were enriched in the significantly upregulated genes of 5% CO_2_ biofilms (NES = +2.62) (Figure 6B). Likewise, significantly differentially expressed genes possessing the GO term ‘amino acid transport’ were primarily upregulated (Figure 6B). The most highly upregulated of these was *GAP2* which encodes a general amino acid permease ^57^. *GAP2* was in fact the highest upregulated gene of the entire RNA-Seq data set with a 4.60 log2 fold change. The basic amino acid permease genes *CAN1, CAN2*, and *CAN3* were also upregulated (Figure 6B).

Genes associated with carbohydrate transmembrane transport were also enriched in the significantly upregulated genes of 5% CO_2_ biofilms as revealed by GSEA (NES = +1.88) (Figure 6B). Amongst these were 12 genes encoding putative major facilitator superfamily (MFS) glucose transmembrane transporters present in the significantly differentially expressed gene list and these are almost all upregulated. The only exceptions are *HGT9* and *HGT18* with log2 fold changes of -1.24 and -1.70 respectively (Figure 6B).

Genes previously identified to be involved in hyphal formation in response to foetal bovine serum (FBS) exposure or 37°C were enriched in the downregulated genes (NES = -3.41). Likewise, the majority of genes under the GO term ‘hyphal growth’ in the significantly differentially expressed gene list were downregulated in 5% CO_2_ biofilms at 48h growth (Supplementary Figure S5B). Significantly differentially expressed genes associated with the cytoskeleton, as identified by GO term analysis, were enriched in the downregulated genes (NES = -2.69) (Supplementary Figure S5C). Cytoskeleton reorganisation is important for the growth of *C. albicans* hyphal cells ^58^ as well as cell division ^59,60^, indicating cell growth is lower in 48h old *C. albicans* biofilms grown in 5% CO_2_ than in those grown in 0.03% CO_2_. Consistent with this, genes involved in the transition through the G1/S checkpoint were also enriched in the significantly downregulated genes (NES = -2.83) (Supplementary Figure S5C).

Initially, these data appear to be contradictory to the previous data highlighting the increased biofilm formation of *C. albicans* under high CO_2_ conditions. However, due to the increased biofilm formation in a 5% CO_2_ environment, *C. albicans* biofilms reach full maturity much quicker in high CO_2_, as observed via confocal microscopy (Figure 2B). Thus, by 48h *C. albicans* biofilms grown in high CO_2_ have been fully mature for several hours, and hence would contain fewer dividing cells or cells extending hyphae in comparison to low CO_2_ biofilms.

### CO_2_ elevation enhances azole resistance in *C. albicans* biofilms

In addition to the observed increase in expression of genes that drive biofilm formation we observed an elevation in certain stress response pathways in *C. albicans* biofilms grown in 5% CO_2_. Gene sets involved in the response of *C. albicans* to antifungals such as Ketoconazole ^61^ (Figure 7A) were upregulated as well as several drug transporters (Figure 7B), indicating that elevation may lead to increased drug resistance. Upregulated genes included the *MDR1* gene (2.56 log2 fold change) which encodes the multidrug resistance pump Mdr1 and is associated with resistance to several antifungals such as azoles ^62^. To test the significance of this *C. albicans* biofilms were seeded and grown for 24h in 0.03% and 5% CO_2_ before the addition of antifungals, after which they were grown for an additional 24h in both conditions to observe the effect of drug application. Antifungal concentrations were selected based upon previously reported MIC values for these antifungals against *C. albicans* biofilms ^28^. Overall, Fluconazole and Miconazole treatment led to a significant reduction in biofilm growth in 0.03% CO_2_ (Figures 7C and Supplementary Figure S6). Treatment with Fluconazole and Miconazole also significantly reduced biofilm formation in 5% CO_2_, however, their effects were markedly reduced (Figures 7C and Supplementary Figure S6). This suggests an increased resistance of biofilms grown in 5% CO_2_ to azole treatment. Interestingly, Nystatin was equally as effective against biofilms in either CO_2_ environment (Figure 7D).

**Figure 7:**
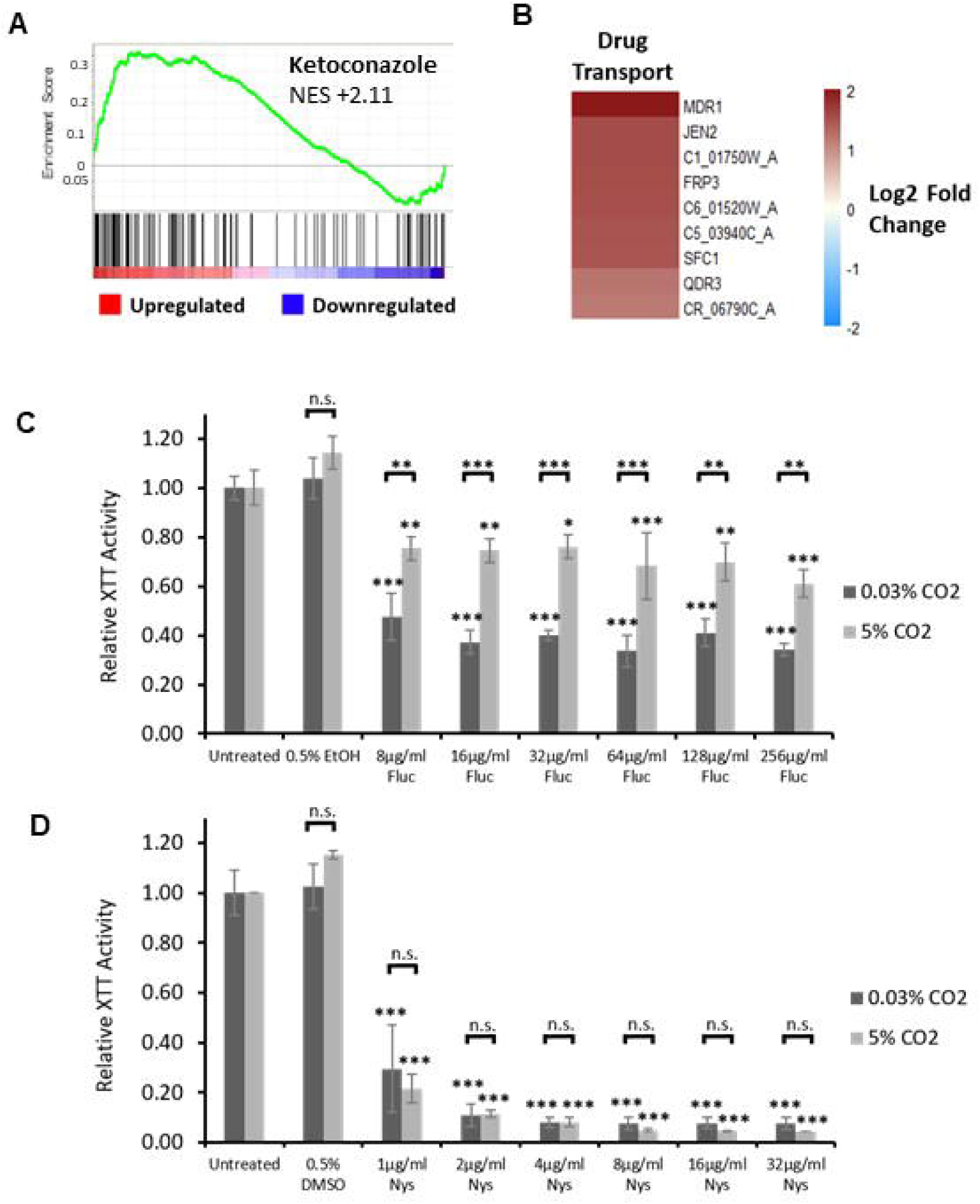
Antifungal sensitivity of *C. albicans* biofilms grown in high (5%) CO_2_. **(A)** GSEA enrichment plot of the KETOCONAZOLE_UP gene set containing genes upregulated in *C. albicans* cells grown in the presence of Ketoconazole ^61^. NES = normalised enrichment score, positive NES indicates enrichment in the upregulated group of genes. **(B)** Heat map of genes associated with drug transport, including the multidrug efflux pump gene *MDR1*. **(C)** Biofilm growth assay of CAI4pSM2 in the presence of Fluconazole. **(D)** Biofilm growth assay of CAI4pSM2 in the presence of Nystatin. The relative XTT activity is presented with the 0.03% CO_2_ biofilms being normalised to the 0.03% CO_2_ untreated control and the 5% CO_2_ biofilms being normalised to the 5% CO_2_ untreated control. Two-way ANOVAs followed by Tukey tests for multiple comparisons were carried out: *p<0.05, **p<0.01, ***p<0.001, n.s. = not significant. Stars directly above the bars indicate a significant difference to untreated in the same CO_2_ environment.

### Precision approaches to overcome CO_2_ acceleration of *C. albicans* biofilm formation

Our data indicate that elevation of CO_2_ leads to an increase in the ability to scavenge for iron and glucose, both essential for biofilm formation and growth. We therefore wished to test whether these represented potential targets to combat *C. albicans* growth in high CO_2_ environments such as the airway. An Fe^3+^ chelator called Deferasirox, which is approved for treating patients with iron overload, has recently been shown to reduce infection levels in a murine oropharyngeal candidiasis model ^63^. With this in mind, we repeated our previous iron starvation biofilm growth assay (Figure 4B) using Deferasirox in place of Ferrozine. We observed that Deferasirox treatment completely eradicates *C. albicans* biofilm growth in 0.03% CO_2_ but has very little effect on biofilm growth in 5% CO_2_ (Figure 8A). Thus adding further evidence that exposure to high levels of CO_2_ can enable *C. albicans* biofilms to overcome the effects of iron starvation. Deferasirox does not therefore appear to be an effective treatment against *C. albicans* biofilms in high CO_2_ such as in the context of voice prostheses colonisation.

**Figure 8:**
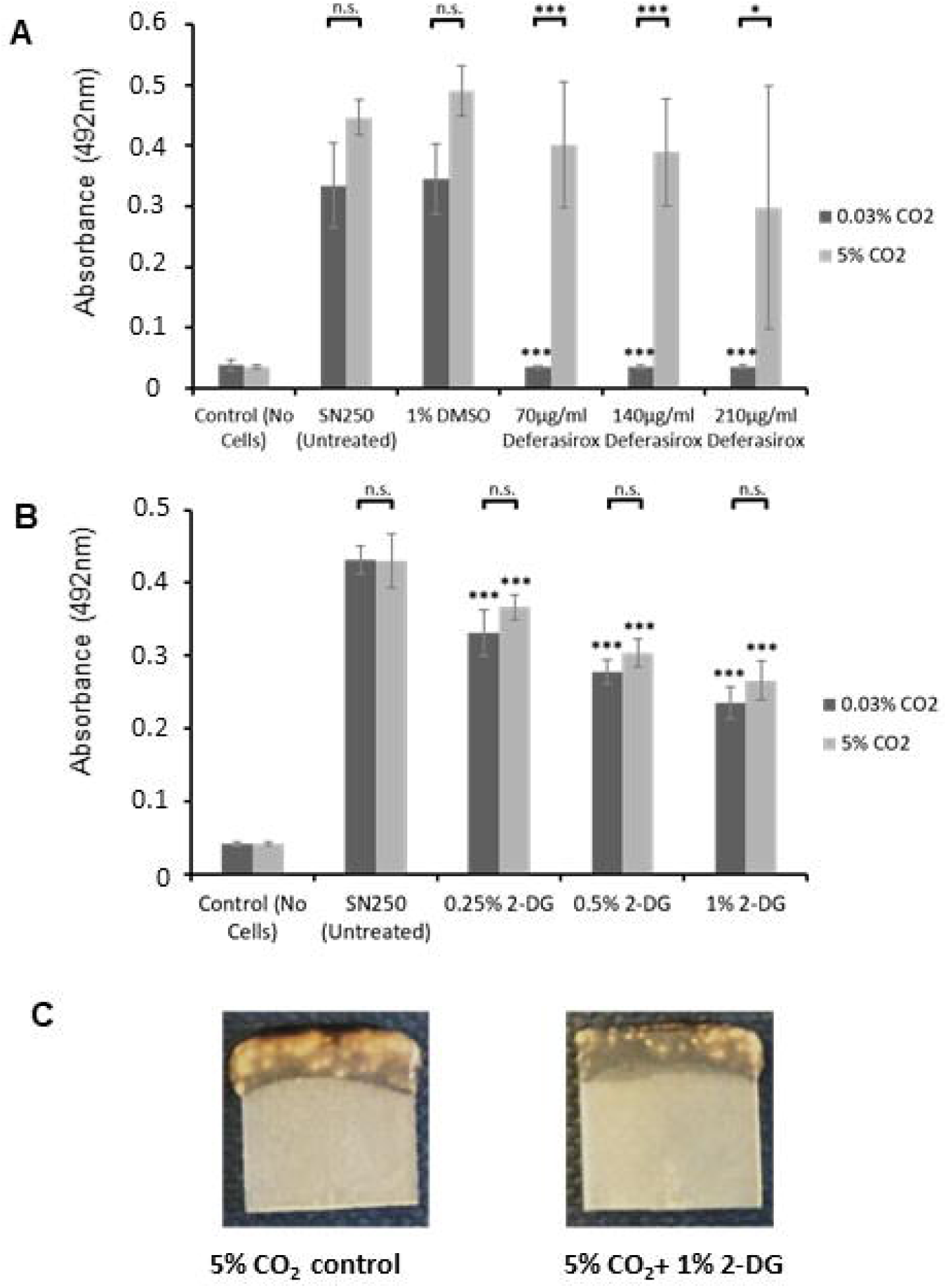
Efficacy of potential treatments to combat *C. albicans* biofilms grown in high (5%) CO_2_. Biofilms were seeded and grown for 48h before XTT quantification. Control wells with no cells were set up as media controls to monitor for contamination. **(A)** Biofilm growth assay of SN250 in the presence of the Fe^3+^ chelator Deferasirox. Graph represents two biological replicates each containing technical triplicates, error bars denote Standard Deviation. **(B)** Biofilm growth assay of SN250 in the presence of the glycolytic inhibitor 2-DG. Graph represents three biological replicates each containing technical triplicates, error bars denote Standard Deviation. Two-way ANOVAs followed by Tukey tests for multiple comparisons were carried out: *p<0.05, **p<0.01, ***p<0.001, n.s. = not significant. Stars directly above the bars indicate a significant difference to the untreated SN250 in the same CO_2_ environment. **(C)** Representative images of SN250 biofilms grown in 5% CO_2_ ±2-DG for 48h.

*C. albicans* biofilms grown in 5% CO_2_ exhibited the upregulation of genes encoding glucose transporters. Accordingly, we contemplated whether treatment of *C. albicans* biofilms with the glucose analogue 2-deoxyglucose (a glycolytic inhibitor) may decrease biofilm growth in high CO_2_ environments. 2-deoxyglucose (2-DG) has previously made it to Stage II clinical trials as an anti-prostate cancer treatment and is considered safe for use in humans ^64^. Thus, it could be a potential therapeutic option to combat *C. albicans* biofilm formation on medical devices, specifically on voice prostheses. *C. albicans* biofilm formation was significantly reduced in the presence of 2-DG regardless of CO_2_ environment (Figure 8B and Supplementary Figure S7). Interestingly, the biofilm reductions in all 2-DG concentrations were similar for biofilms in both low and high CO_2_, despite the fact several glucose transporters were upregulated in 5% CO_2_ biofilms (Figure 6B). The reduction in biofilm growth upon 2-DG treatment was also quite apparent macroscopically (Figure 8C).

## Discussion

Our data demonstrate for the first time how a physiologically relevant elevation of CO_2_ accelerates biofilm formation in *C. albicans* by activating the cAMP/PKA pathway (Figure 9). Although CO_2_ elevation is dependent on Cyr1 it appears to bypass a requirement for Ras1. CO_2_ elevation enhances each stage of the *C. albicans* biofilm forming process, from attachment through maturation to dispersion. The observed increase in cell attachment is accompanied by an increase in the abundance of mRNA transcripts for *ALS1, ALS2* and *ALS4* which encode adhesins of the agglutinin-like sequence family that function in the cell-surface and cell-cell attachment of *C. albicans* ^19^. Our observed increase in dispersion also correlates with an upregulation of the known regulator of this stage of biofilm growth, *NRG1* ^65^. Our observations have important clinical implication in scenarios where prosthetic devices are placed in areas of elevated CO_2_, for example voice prostheses or tracheostomy tubing. We would anticipate that high CO_2_ may increase the probability of *C. albicans* colonisation and dissemination.

**Figure 9:**
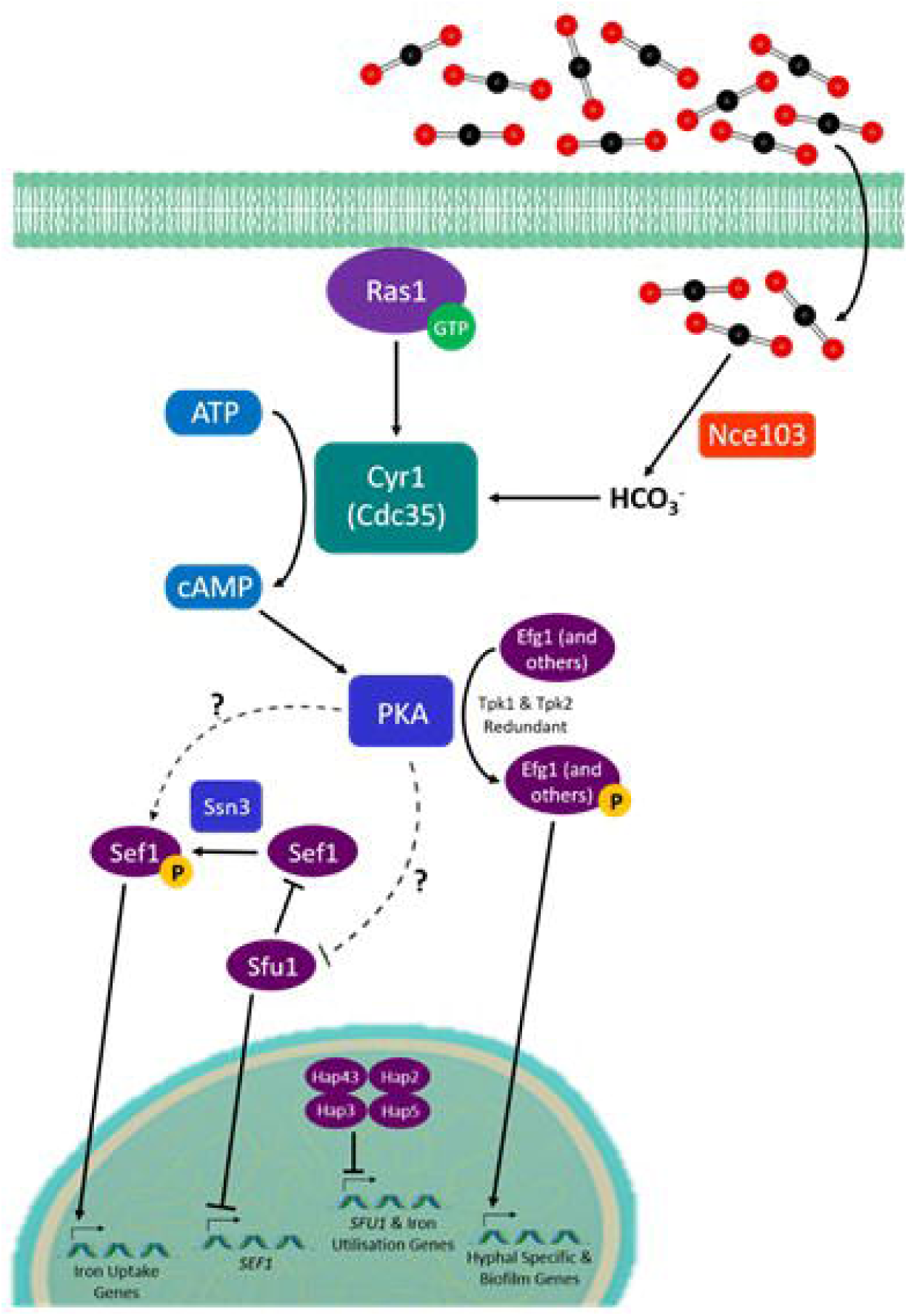
Predicted model of the interplay between CO_2_ signalling and iron homeostasis in *C. albicans* biofilms on silicone surfaces. We hypothesise that the CO_2_-mediated activation of PKA via Cyr1 will result in increased activity of Sef1 (via its phosphorylation and possibly the inhibition of Sfu1), thus increasing the expression of iron-uptake genes. PKA activation leads to Efg1 activation promoting the biofilm process.

Of the original six ‘core’ biofilm regulators (Efg1, Bcr1, Brg1, Rob1, Ndt80 and Tec1) only four (Efg1, Bcr1, Brg1, and Rob1) were identified as required for normal biofilm growth in our TFKO screen. The original TFKO screen which identified these regulators was carried out using a polystyrene surface ^27^ whereas we used silicone. This may suggest that attachment to and biofilm growth on silicone requires a more limited set of core transcription factors reflecting the importance of surface upon biofilm formation as has been observed previously ^55^. In support of the environment being critical to biofilm establishment, CO_2_ elevation was able to compensate for the loss of the core biofilm regulators Brg1 and Rob1. Intriguingly, of the four biofilm regulators whose gene sets were predicted to be upregulated in 5% CO_2_ biofilms, Efg1, Bcr1 and Brg1 were identified in our TFKO screen whereas Ndt80 was not. Moreover, only *efg1Δ/Δ* and *bcr1Δ/Δ* had reduced biofilm growth in both 0.03% and 5% CO_2_. This disparity may be explained by the degree of overlap in downstream target genes between the core regulators of biofilm formation ^27^ which implies significant potential for functional redundancy.

Many of the genes significantly upregulated in biofilms grown in high CO_2_ after 48h were also found to be upregulated in other biofilm gene expression studies ^66,67^. For instance, Nett, J. *et al.* observed an increase in the expression of adherence genes in an *in vivo* venous catheter biofilm model. Like us, this venous catheter biofilm study saw an increase in the transcript abundance of *ALS1* and *ALS2* however, this was only observed at earlier time point biofilms ^66^. This was also the case in a temporal gene expression analysis using *in vitro* denture and catheter models ^67^. Interestingly, some pathways such as hexose transport, amino acid uptake, and stress responses that our differential gene expression analysis predicted to be upregulated in mature biofilms grown in high CO_2_ were concluded to be upregulated only in early phase biofilms (12h) in this denture and catheter study ^67^. The authors concluded the result of the induction of these pathways is the increase of intracellular pools of pyruvate, pentoses and amino acids, preparing for the large increase in biomass that occurs later in biofilm development ^67^. We hypothesise that high CO_2_ may stimulate these pathways and maintain their activity even in mature biofilms, thus supporting the increased biomass and maturation rate of biofilms observed when grown in high CO_2_.

We observed an increase in azole antifungal resistance within *C. albicans* biofilms grown in 5% CO_2_. This, at least partly, could be explained by the increase in expression of drug transporter genes such as *MDR1* which have previously been implicated in azole resistance ^62,68^. However, *mdr1Δ/Δ*, as well as *cdr1Δ/Δ* and *cdr2Δ/Δ*, mutants only exhibit reduced azole resistance in planktonic culture and early stage (6h) biofilms, while levels of resistance are maintained in mature biofilms ^62^. Therefore, we propose that it is more likely the increased azole resistance phenotype of 5% CO_2_ biofilms displayed is contributed to via another mechanism, possibly increased ECM deposition. β-1,3-glucan, a major component of biofilm ECM, can bind to azole antifungals and sequester them to prevent passage to the cells ^69^. We have observed that after 48h, 5% CO_2_ biofilms, while containing similar cell numbers to 0.03% CO_2_ biofilms, often appear larger to the eye with a more bulbous appearance. This could suggest more ECM material being produced in high CO_2_ environments, contributing to the increased azole resistance. Furthermore, Miconazole treatment has been shown to generate superoxide radicals within *C. albicans* biofilms and leads to the increased expression of *SOD5* and *SOD6* (encode superoxide dismutase enzymes) in an attempt to protect against the toxic superoxides. A *sod4Δ/Δsod5Δ/Δsod6Δ/Δ* triple mutant is hypersensitive to Miconazole treatment when growing as a biofilm ^70^. Our transcriptome analysis revealed *SOD6* and *SOD4* are upregulated in 5% CO_2_ biofilms, providing a potential further mechanism for increased Miconazole resistance.

Our study identified that transcription factors involved in iron homeostasis are important for *C. albicans* biofilm growth. Principal among these were the Hap transcription factors which come together to form the HAP complex, a CCAAT box-binding transcriptional regulator, under iron-limiting conditions. Genetic studies have revealed a requirement of *HAP2, HAP3, HAP5* and *HAP43* for growth in low-iron media ^38,71^. Thus, the biofilm formation defect exhibited by the *hap2Δ/Δ, hap3Δ/Δ, hap5Δ/Δ*, and *hap43Δ/Δ* mutants in 0.03% CO_2_ could be explained by this growth deficiency since RPMI-1640 media has a low iron content. Nevertheless, this makes it even more intriguing that simply an increase in ambient CO_2_ levels was able to significantly increase the biofilm growth of these mutants.

The HAP complex represses a GATA-type transcription factor called Sfu1. Sfu1 is responsible for repressing iron-uptake genes along with *SEF1* under iron-replete conditions. Sef1 activates iron-uptake genes as well as *HAP43, HAP2* and *HAP3* ^52^, in this way the HAP complex is able to induce iron-uptake pathways while repressing iron-utilisation genes ^51,52^. Deletion of Sef1 results in the aberrant downregulation of all the major iron-uptake pathways of *C. albicans* in low iron conditions ^52^. Due to the fact the *sef1Δ/Δ* mutant had defective biofilm growth in both CO_2_ conditions, we hypothesise that a high CO_2_ environment may influence Sef1 directly, causing the HAP complex to become at least partially redundant under these conditions. It should be noted that we did not observe a significant effect on *SEF1* mRNA levels within 5% CO_2_ biofilms after 48h. Thus, if it is influencing Sef1 activity, CO_2_ may be acting at either the protein level or post-translational level as Sef1 is subject to a post-translational control loop consisting of Sfu1 and the cyclin-dependent kinase Ssn3 ^72^. However Ssn3 may not play a role as it has recently been found to be dephosphorylated, and thus inactive, in 5% CO_2_ ^73^. Our current model suggests that under elevated CO_2_ PKA can phosphorylate Sef1 and possibly also inhibit Sfu1 (Figure 9). This is currently under investigation and may possibly explain the tolerance to iron sequestration exhibited by *C. albicans* biofilms grown in high CO_2_. We believe any potential phosphorylation of Sef1 by PKA to be Tpk1/Tpk2 redundant since the *tpk1Δ/Δ* and the *tpk2Δ/Δ* mutants both exhibited a significant increase in biofilm growth in 5% CO_2_ when in the presence of 500*µ*M Ferrozine compared to 0.03% CO_2_. This is supported by a Tpk1/Tpk2 phosphoproteomic study which predicted Sef1 to be a potential PKA target that can be phosphorylated by both Tpk1 and Tpk2 ^74^.

The increased azole resistance of *C. albicans* biofilms in 5% CO_2_ along with the observed iron starvation tolerance when treated with the Fe^3+^ chelator Deferasirox has important implications for the development of potential biofilm treatment strategies. At a clinical level this also indicates the location of a *C. albicans* biofilm within the body should be taken into consideration when deciding upon the most effective treatment. Encouragingly, 2-deoxyglucose (2-DG) was able to attenuate *C. albicans* biofilm growth in both 5% and atmospheric (0.03%) CO_2_ environments. 2-DG has exhibited antimicrobial effects against fungal moulds ^75^ and bacterial biofilms ^76^. This, together with its action against *C. albicans* biofilms presented here, highlights the potential for 2-DG to be used an anti-biofilm therapeutic. It may be particularly useful for medical devices such as voice prostheses which are situated in CO_2_-rich environments in the body and are often colonised by a mixture of bacterial and fungal species ^77,78,79,80^. It’s important to note however that 2-DG was unable to eradicate *C. albicans* biofilm growth completely. It may be that 2-DG treatment is beneficial in combination with other compounds, such as iron chelators or traditional antifungals; this possibility is yet to be explored.

In conclusion, our data demonstrates that elevated levels of CO_2_ act as an important driver of *C. albicans* biofilm formation, growth and maturation. These CO_2_-mediated effects are likely to have important medical ramifications, particularly in the context of prosthetics and airway management devices, but also for host infections in CO_2_-rich environments in the body.

## Acknowledgments

Our gratitude is extended to the Kent Cancer Trust who provided funds to support Mr. Daniel Pentland through his PhD and to the contributions of the East Kent Hospital University Foundation Trust in integration of this research within new *Candida* management voice prosthesis pathway guidelines.

## Author Contributions

The study was conceived by C.W. Gourlay and F.A. Mühlschlegel. Experimental procedures and data analyses were conducted by D.R. Pentland. The manuscript was written and edited by D.R. Pentland, C.W. Gourlay and F.A. Mühlschlegel.

