## Supplemental tables and figures for "CO_2_ enhances the ability of *Candida albicans* to form biofilms, overcome nutritional immunity and resist antifungal treatment"

**Supplementary Table S1**

| Strain | Species | Genotype/Parent | Source/Reference |
|---|---|---|---|
| SN152 | *C. albicans* | *ura3/*::imm434::*URA3/ura3*::imm434 *iro1::IRO1/iro1*::imm434 *his*1::hisG/*his1*::hisG *leu2/leu2 arg4/arg4* | [39] |
| SN250 (TFKO Library Reference Strain) | *C. albicans* | *ura3/*::imm434::*URA3/ura3*::imm434 *iro1::IRO1/iro1*::imm434 *his*1::hisG/*his1*::hisG *leu2*::*CdHIS1/leu2*::*CmLEU2 arg4/arg4* | [39] |
| CAI4 | *C. albicans* | *ura3*::imm434/*ura3*::imm434 *iro1/iro1*::imm434 | Mühlschlegel Lab |
| CAI4pSM2 | *C. albicans* | CAI4 transformed with pSM2, *URA3* integrating plasmid | [66] |
| TFKO Library | *C. albicans* | SN152 | [39] |
| CDH107 (*ras1Δ/Δ*) | *C. albicans* | CAI4 | [32] |
| CR276 (*cdc35Δ/Δ*) | *C. albicans* | CAI4 | [84] |
| WYF2 ($CDC35^{\Delta RA}$) | *C. albicans* | CR276 transformed with pClp-$cdc35^{\Delta RA}$ | [69] |
| *tpk1Δ/Δ* | *C. albicans* | CAI4 | Mühlschlegel Lab |
| *tpk2Δ/Δ* | *C. albicans* | CAI4 | Mühlschlegel Lab |
| G-3065 | *C. albicans* | Clinical Isolate | Failed Voice Prosthesis |
| G-8424 | *C. albicans* | Clinical Isolate | Failed Voice Prosthesis |
| G-1625 | *C. albicans* | Clinical Isolate | Failed Voice Prosthesis |

**Supplementary Table S1: *Candida albicans* strains used in this study.**

**Supplementary Table S2**

| TFKO Mutant | Biofilm Growth | |
|---|---|---|
| | 0.03% $CO_2$ | 5% $CO_2$ |
| *tup1Δ/Δ* | – – – (p < 0.001) | – – – (p < 0.001) |
| *sef1Δ/Δ* | – – (p = 0.002) | – – (p < 0.001) |
| *swi4Δ/Δ* | – – (p < 0.001) | – – (p = 0.002) |
| *pho4Δ/Δ* | – – – (p < 0.001) | – (p = 0.049) |
| *bcr1Δ/Δ* | – – – (p < 0.001) | – – (p = 0.009) |
| *efg1Δ/Δ* | – – (p = 0.007) | – – – (p = 0.044) |
| *hap2Δ/Δ* | – – – (p < 0.001) | n.s. (p = 0.561) |
| *rbf1Δ/Δ* | – – (p < 0.001) | n.s. (p = 0.873) |
| *rob1Δ/Δ* | – – (p < 0.001) | n.s. (p = 0.477) |
| *fgr15Δ/Δ* | – – (p = 0.005) | n.s. (p = 0.637) |
| *dal81Δ/Δ* | – – (p = 0.007) | n.s. (p = 0.682) |
| *mig1Δ/Δ* | – – (p = 0.014) | n.s. (p = 0.264) |
| *brg1Δ/Δ* | – – (p = 0.022) | n.s. (p = 0.998) |
| *C4_00260WΔ/Δ* | – – (p = 0.023) | n.s. (p = 0.915) |
| *zcf27Δ/Δ* | – – (p = 0.047) | n.s. (p = 0.870) |
| *C1_13880CΔ/Δ* | – (p = 0.009) | n.s. (p = 0.803) |
| *crz1Δ/Δ* | – (p = 0.022) | n.s. (p = 0.474) |
| *hap43Δ/Δ* | – (p = 0.030) | n.s. (p = 0.338) |
| *leu3Δ/Δ* | n.s. (p = 0.120) | – – (p < 0.001) |
| *mbp1Δ/Δ* | n.s. (p = 0.085) | – – (p = 0.007) |
| *bas1Δ/Δ* | n.s. (p = 0.563) | – (p = 0.016) |
| *try6Δ/Δ* | n.s. (p = 1.000) | – (p = 0.037) |
| *mac1Δ/Δ* | ++ (p = 0.020) | n.s. (p = 0.847) |
| *zcf30Δ/Δ* | ++ (p = 0.038) | n.s. (p = 0.640) |
| *zcf17Δ/Δ* | + (p = 0.023) | n.s. (p = 0.735) |

**Supplementary Table S2: Summary of the transcription factor knockout (TFKO) mutants which had significantly altered biofilm growth in 0.03% and/or 5% $CO_2$ environments within our screen.** Biofilms were quantified via XTT assays and normalised to the wild type XTT readout within each $CO_2$ environment; – – – <40% XTT activity relative to wild type, – – 40-70%, – >70%, + 110-130%, ++ >130%, n.s. non-significant. Green indicates transcription factors previously known to have a role in biofilm regulation as per Gene Ontology analysis. Grey indicates transcription factors involved in maintaining cellular iron homeostasis.

**Supplementary Figure S1**

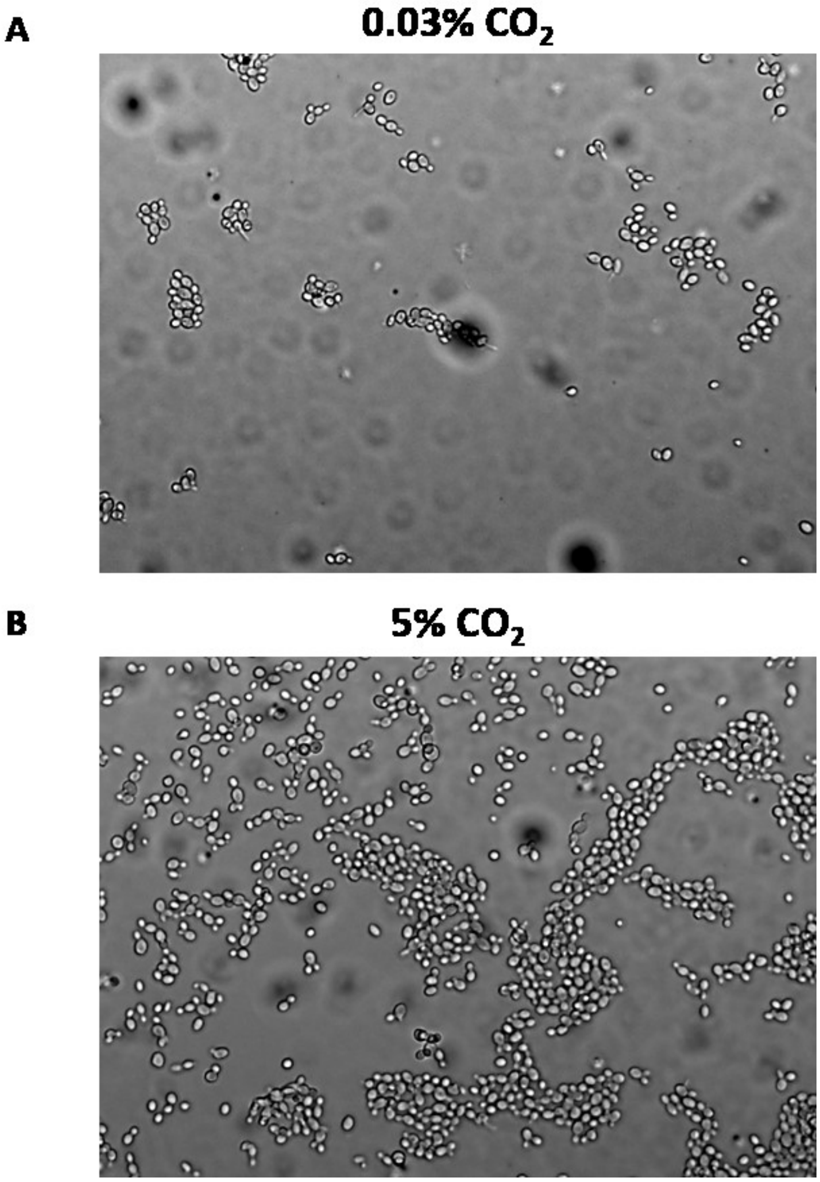


**Supplementary Figure S1: Attachment of *C. albicans* CAI-4 cells to a silicone surface in 0.03% and 5% $CO_2$.** *C. albicans* CAI-4 cells were seeded for 90 mins onto silicone-coated microscope slides in 0.03% $CO_2$ or 5% $CO_2$, unattached cells were washed off and images taken at 20x objective magnification. Several images were taken of both 0.03% and 5% $CO_2$ slides and representative examples are shown. **(A)** Representative image of a slide section from 0.03% $CO_2$. **(B)** Representative image of a slide section from 5% $CO_2$.

**Supplementary Figure S2**

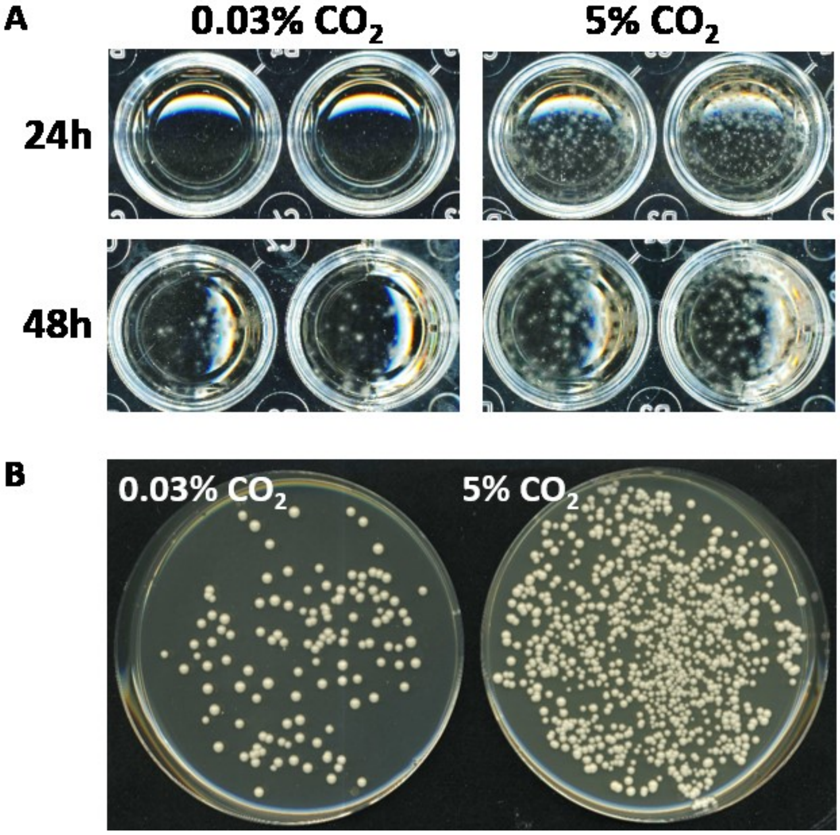


**Supplementary Figure S2: Dispersion of *C. albicans* CAI4pSM2 cells from biofilms grown in 0.03% and 5% $CO_2$. (A)** Wells containing RPMI-1640 media after biofilm growth, *C. albicans* cells can be seen as white clumps formed from hyphal cells. **(B)** Representative CFU plates from a 1:10 dilution of the spent RPMI-1640 media after 48h biofilm growth in both $CO_2$ conditions.

**Supplementary Figure S3**

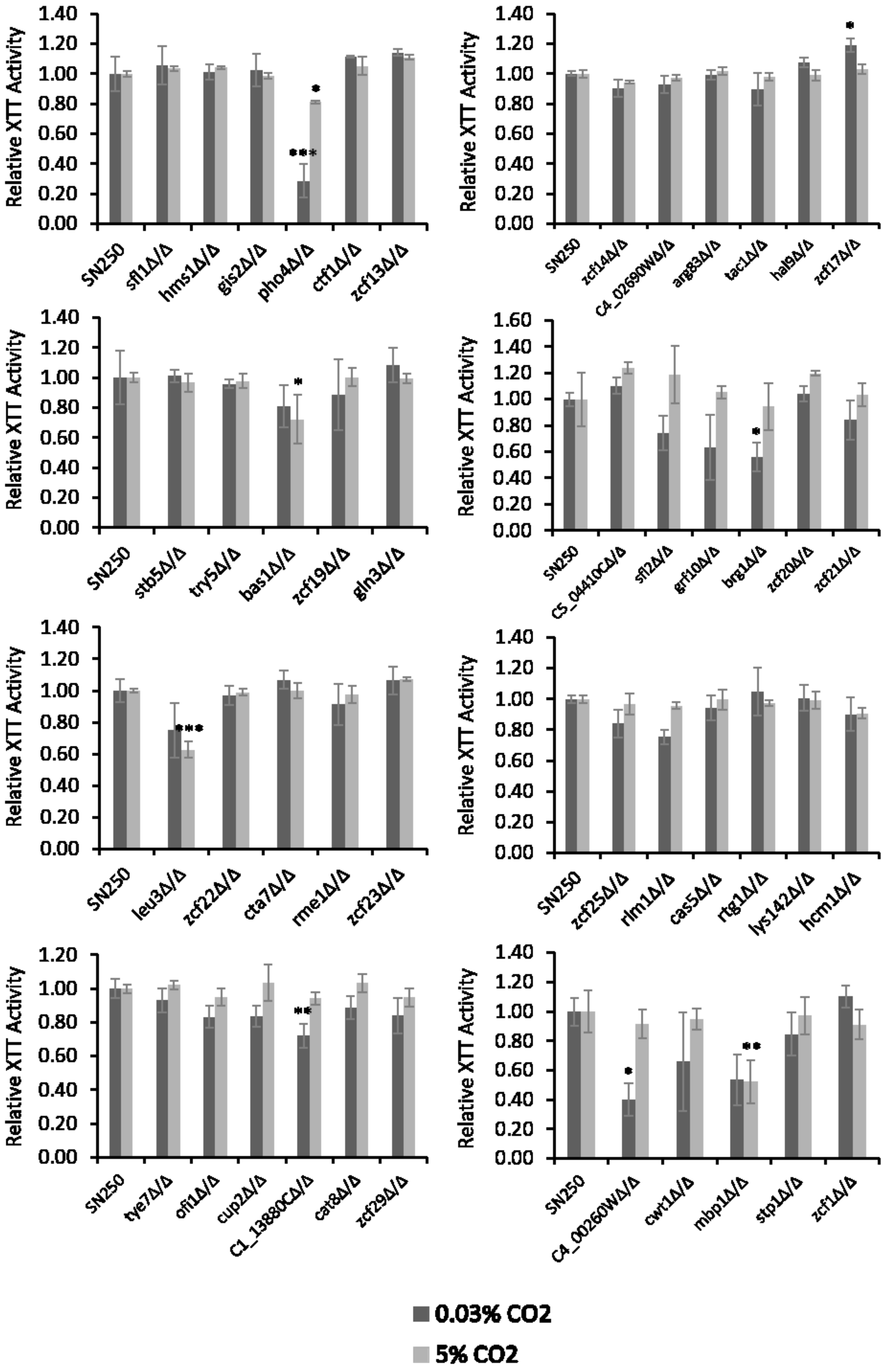

**Supplementary Figure S3**

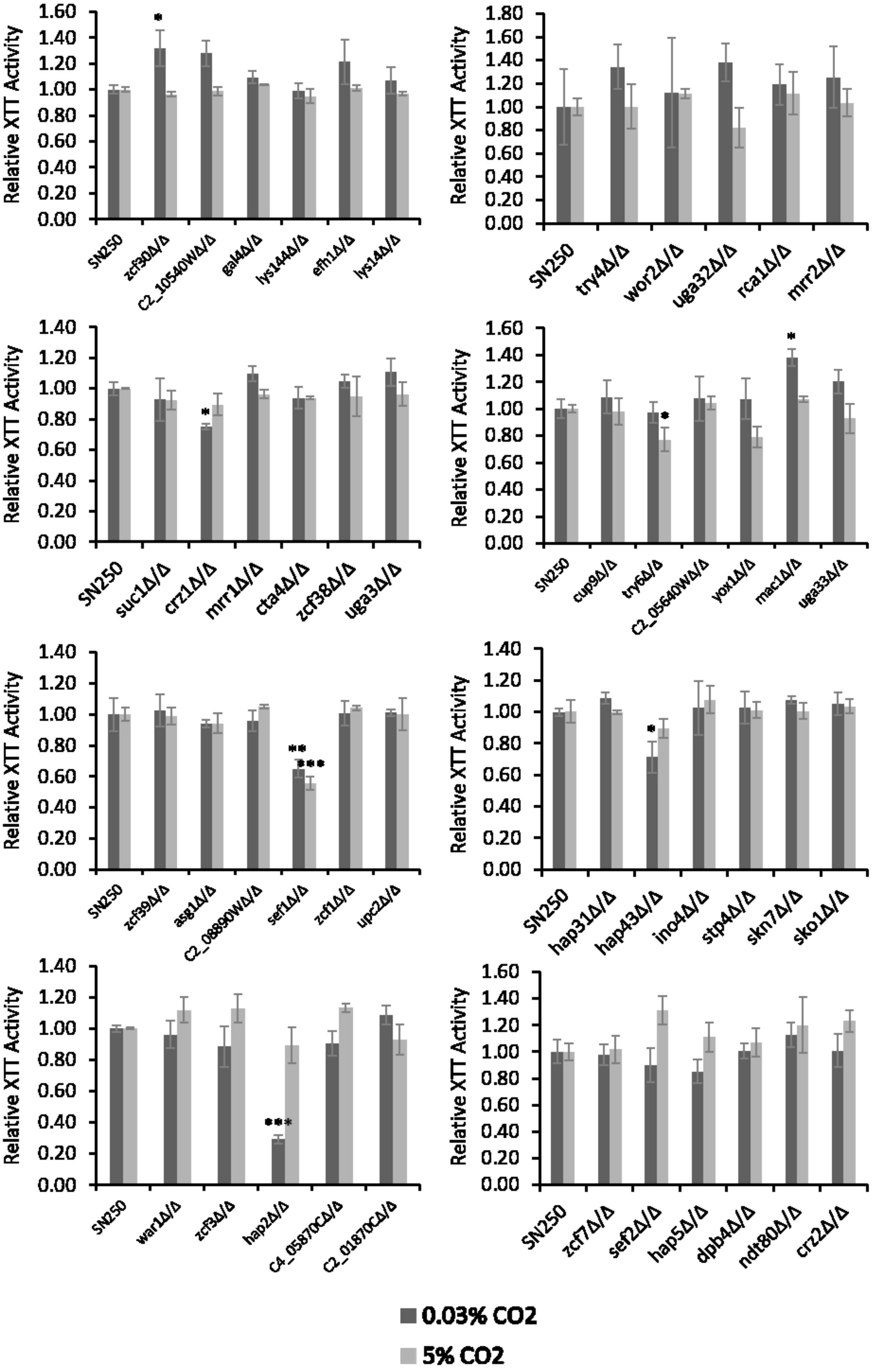

**Supplementary Figure S3**

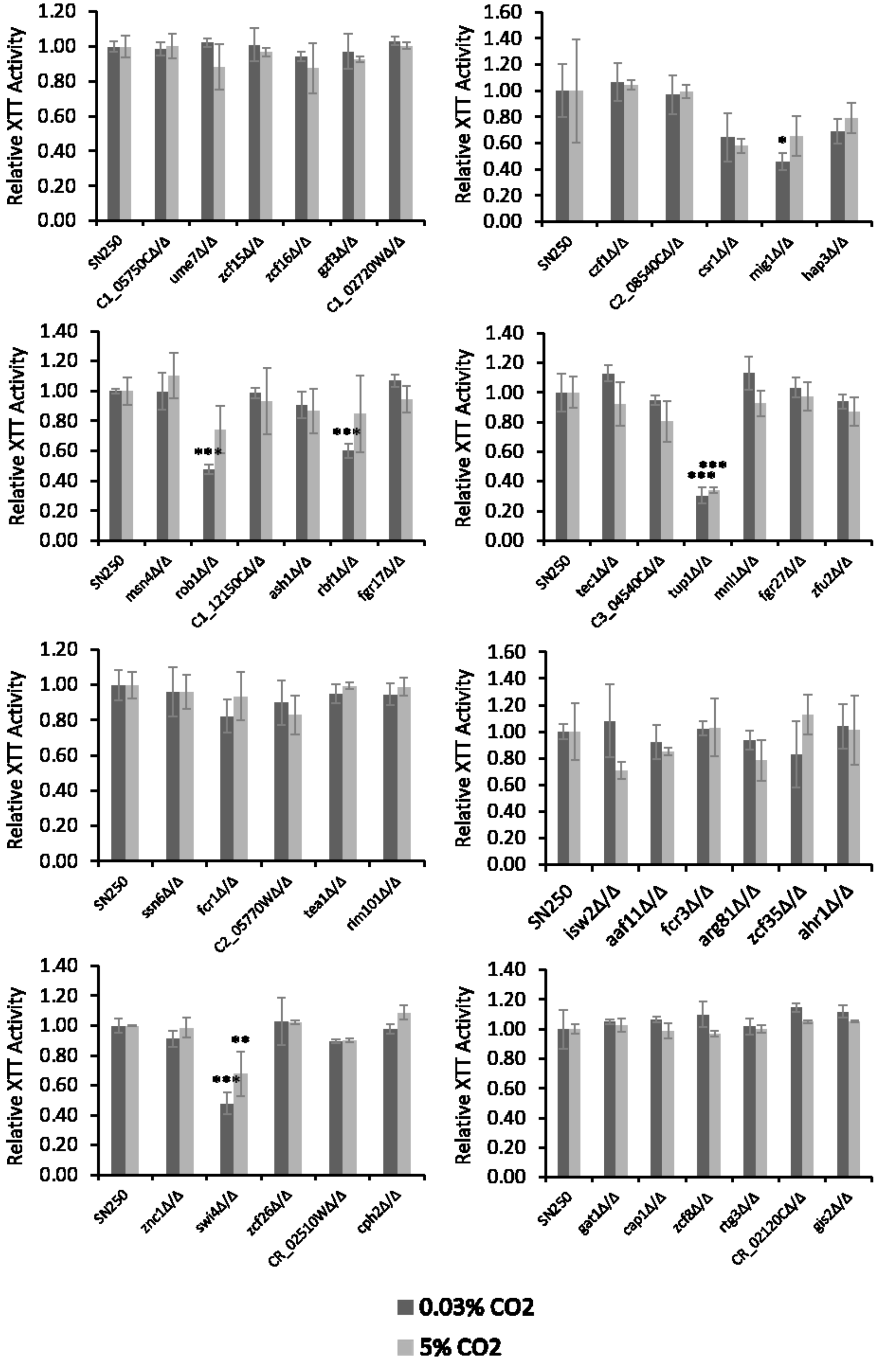

**Supplementary Figure S3**

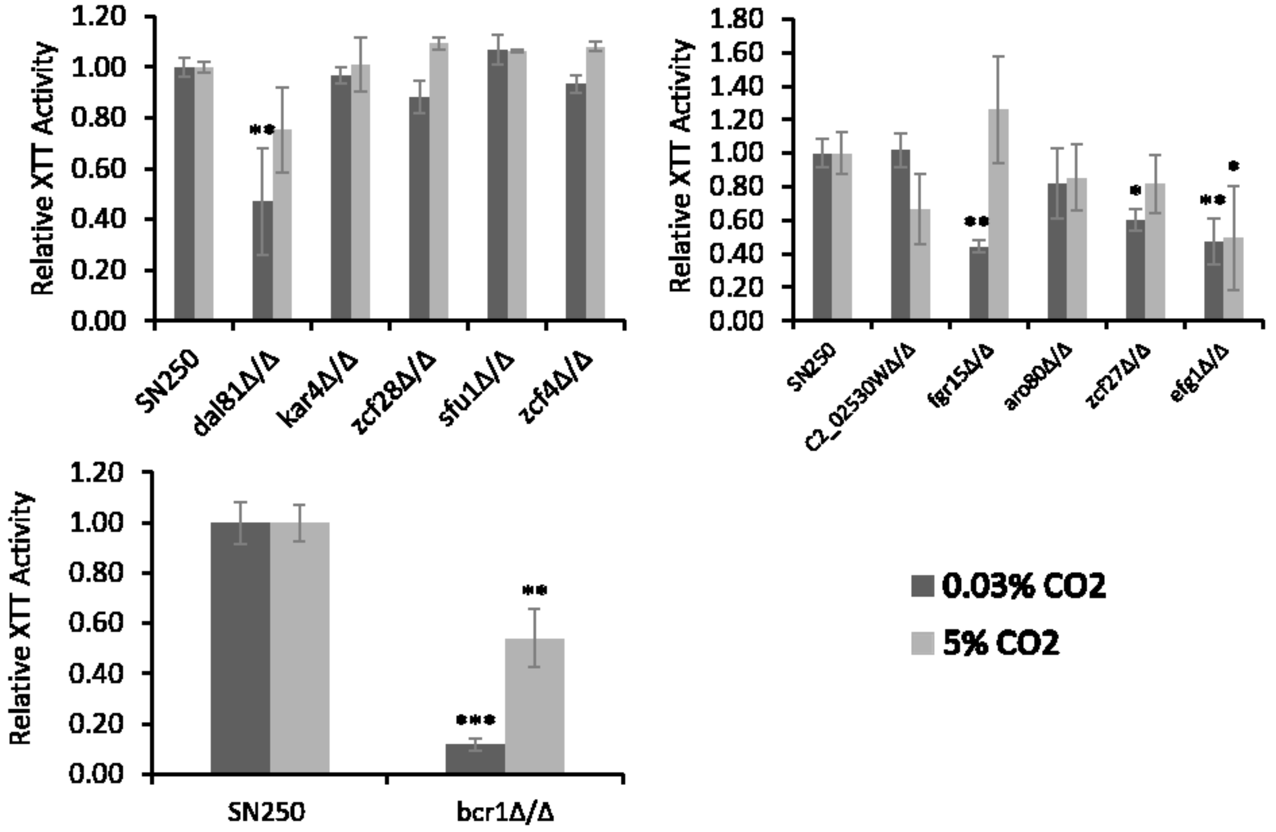


**Supplementary Figure S3: Transcription Factor Knockout Screen of *C. albicans* biofilm-forming ability.** Biofilms were seeded and grown before quantification via XTT assay. Graphs represent three biological replicates for each mutant. The XTT assay absorbances at 492nm for the 0.03% and 5% $CO_2$ biofilms have been normalised to the 0.03% and 5% $CO_2$ SN250 wild type controls respectively. This removes any day-to-day variation, thus allowing TFKO mutants grown on different days to be compared. One-way ANOVAs followed by Dunnett's Tests for multiple comparisons to a control were performed to compare the TFKO mutant biofilms to the SN250 wild type controls (for both 0.03% and 5% $CO_2$); *p<0.05, **p<0.01, ***p<0.001.

**Supplementary Figure S4**

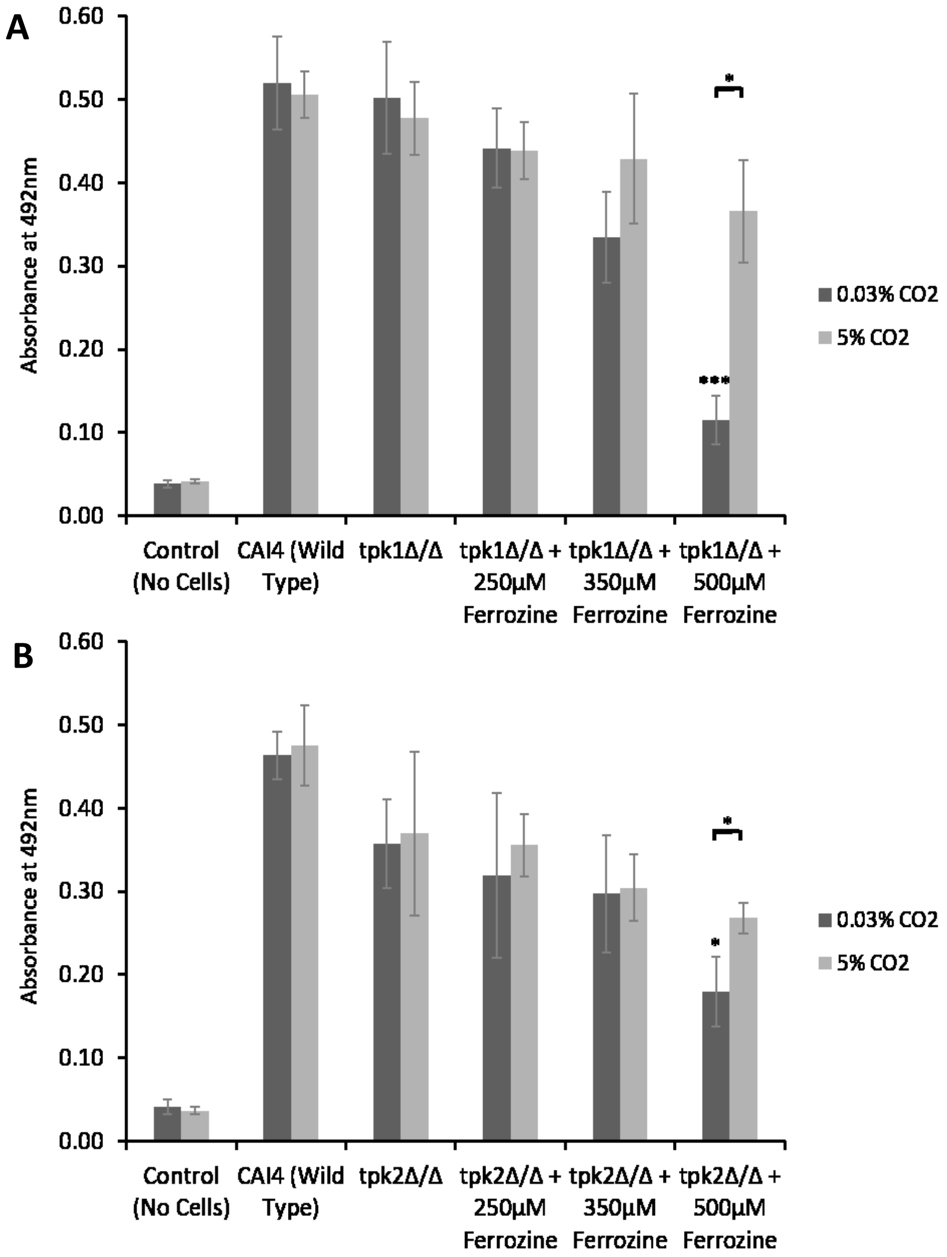


**Supplementary Figure S4: The effect of high (5%) $CO_2$ on tolerance to iron starvation in *tpk1Δ/Δ* and *tpk2Δ/Δ* PKA mutants.** Biofilms using **(A)** *tpk1Δ/Δ* and **(B)** *tpk2Δ/Δ* mutants were seeded and grown for 48h in the presence of the $Fe^{2+}$ chelator Ferrozine before XTT quantification. Graphs represents two biological replicates each containing technical triplicates, error bars denote Standard Deviation. Two-way ANOVAs followed by Tukey tests for multiple comparisons were carried out: *p<0.05, **p<0.01, ***p<0.001, n.s. = not significant. Stars directly above the bars indicate a significant difference to the untreated *tpk* deletion strain in the same $CO_2$ environment.

**Supplementary Figure S5**

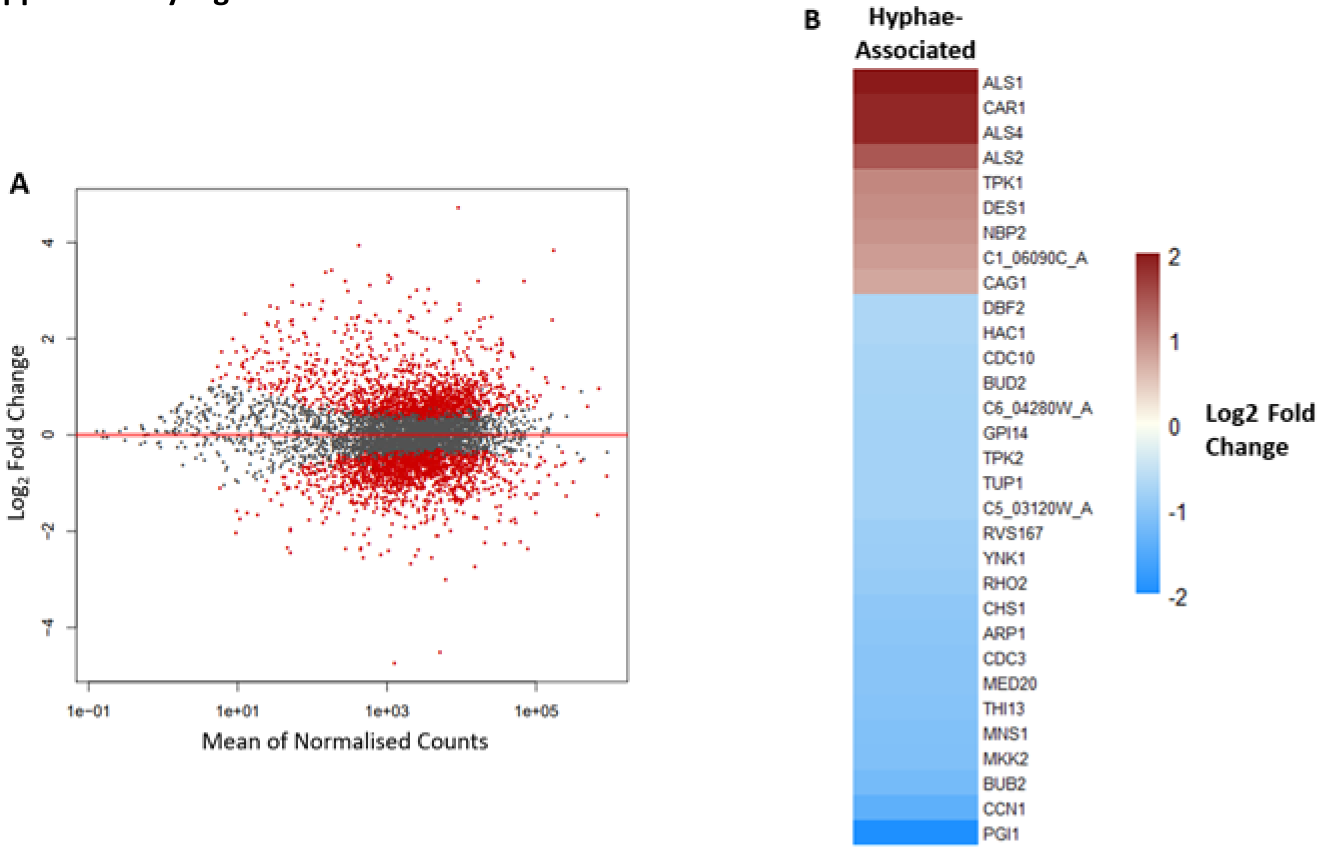


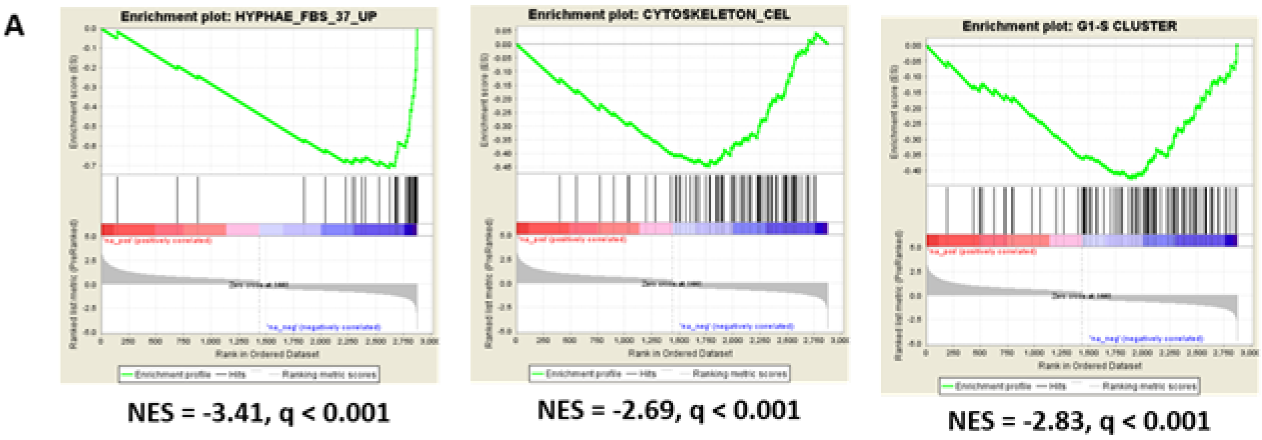


**Supplementary Figure S5: Gene expression profiles of genes/gene sets downregulated in 5% $CO_2$ vs. 0.03% $CO_2$ *C. albicans* biofilms. . (A)** Volcano plot of $\log_2$ fold changes (5% $CO_2$ vs. 0.03% $CO_2$ biofilms) against mean of normalised counts for each gene; red dots = significantly differentially expressed (q≤0.05) genes, grey dots = not significant. 2875 genes showed significant differential expression. Normalised counts are the number of reads a particular gene has, thus the higher the mean of normalised count, the higher the expression of that gene.**(A)** GSEA enrichment plots of the HYPHAE_FBS_37_UP gene set containing genes upregulated after 6h of exposure to FBS and 37°C (hyphal inducing conditions), the CYTOSKELETON_CEL gene set containing genes under the GO term 'cytoskeleton', and the G1-S CLUSTER gene set containing genes involved in the transition through the G1/S checkpoint. Vertical black lines represent individual genes in the ranked gene list from upregulated (left) to downregulated (right). NES = normalised enrichment score, negative NES indicates enrichment in the downregulated group of genes. **(B)** Heatmap of genes associated with hyphal growth as identified by GO term analysis. Colours saturate at log2 fold change of 2 and -2; *ALS1* actually has a log2 fold change of 3.77.

**Supplementary Figure S6**

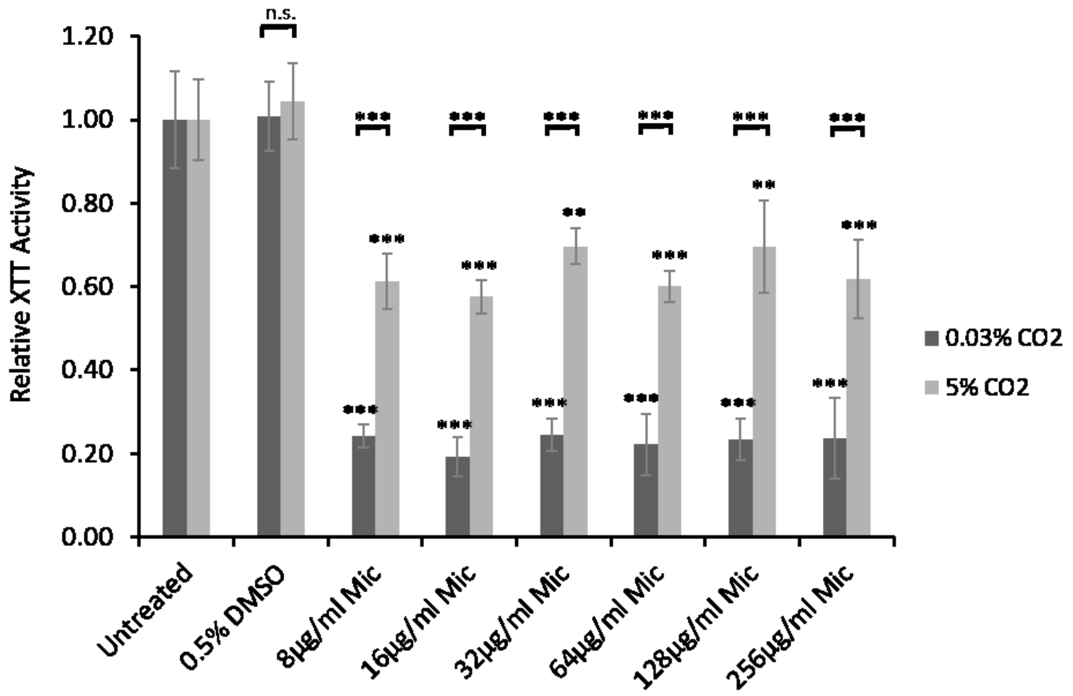


**Supplementary Figure S6: Miconazole sensitivity of *C. albicans* biofilms grown in high (5%) $CO_2$.** Biofilm growth assay of CAI4pSM2 in the presence of Miconazole. Biofilms were seeded and grown for 24 hours before antifungal addition, they were then grown for a further 24 hours before quantification using the XTT assay. The relative XTT activity is presented with the 0.03% $CO_2$ biofilms being normalised to the 0.03% $CO_2$ untreated control and the 5% $CO_2$ biofilms being normalised to the 5% $CO_2$ untreated control. This prevents the general higher growth of 5% $CO_2$ biofilms impacting the analysis. Two-way ANOVAs followed by Tukey tests for multiple comparisons were carried out: *p<0.05, **p<0.01, ***p<0.001, n.s. = not significant. Stars directly above the bars indicate a significant difference to untreated in the same $CO_2$ environment.

**Supplementary Figure S7**

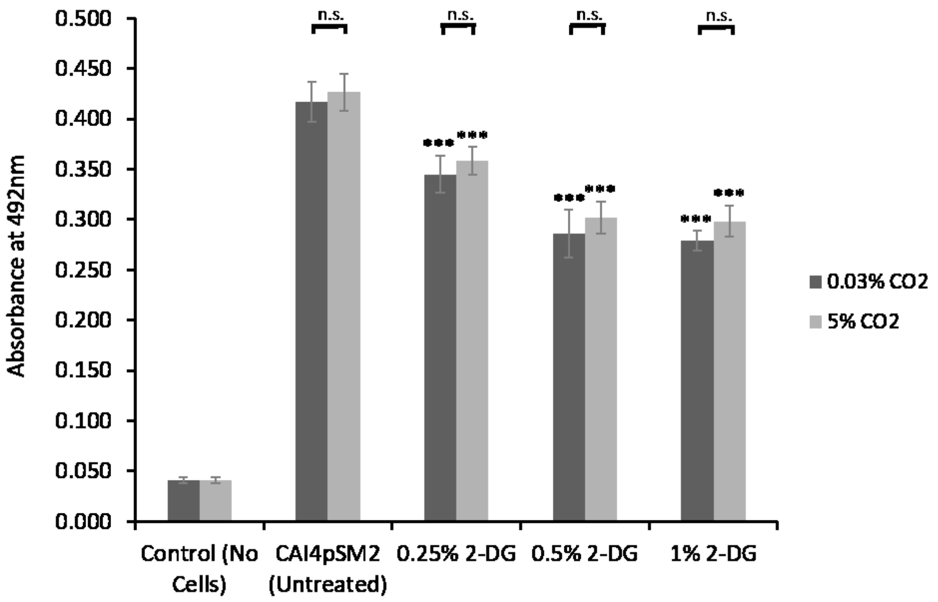


**Supplementary Figure S7: Treatment of CAI4pSM2 *C. albicans* biofilms with 2-DG in 0.03% and 5% $CO_2$.** Biofilms were seeded and grown for 48h in the presence of the glycolytic inhibitor 2-DG before XTT quantification. Control wells with no cells were set up as media controls to monitor for contamination as well as to ensure there was no reaction of the silicone squares with the XTT reagents. Graph represents three biological replicates each containing technical triplicates, error bars denote Standard Deviation. Two-way ANOVAs followed by Tukey tests for multiple comparisons were carried out: *$p<0.05$, **$p<0.01$, ***$p<0.001$, n.s. = not significant. Stars directly above the bars indicate a significant difference to the untreated CAI4pSM2 in the same $CO_2$ environment.
